# Disease-specific tau polymorphs define unique protein interaction networks across proteinopathies

**DOI:** 10.1101/2025.09.09.675096

**Authors:** Nicha Puangmalai, Nemil Bhatt, Nopparat Suthprasertporn, Ernesto G. Miranda-Morales, Nikita Shchankin, Madison Samples, Ahmet E. Balci, Jia Yi Liew, Mauro Montalbano, Yingxin Zhao, Rakez Kayed

**Affiliations:** Mitchell Center for Neurodegenerative Diseases, University of Texas Medical Branch, Galveston, Texas, USA; Departments of Neurology, Neuroscience and Cell Biology, University of Texas Medical Branch, Galveston, Texas, USA; Department of Internal Medicine, University of Texas Medical Branch, Galveston, Texas, USA; Institute for Translational Sciences, University of Texas Medical Branch, Galveston, TX, 77555, USA

**Keywords:** Tau polymorphs, Tau Interactome, Disease Heterogeneity, Tau Post-Translational Modifications

## Abstract

Tau protein aggregates exhibit distinct conformations across tauopathies, but their disease-specific protein interactions remain poorly understood. Here, we demonstrate that disease-specific tau conformations determine unique protein interaction landscapes across Alzheimer’s disease (AD), progressive supranuclear palsy (PSP), and dementia with Lewy bodies (DLB). Through comprehensive interactome profiling of misfolded tau aggregates from PBS- and sarkosyl-soluble fractions. We identified 493 high-confidence proteins with remarkable disease specificity—notably, no common interactors overlapping across all three tauopathies. Machine learning classification achieved compelling discrimination between diseases using as few as 4-6 proteins features, demonstrating robust molecular signatures underlying clinical heterogeneity. AD derived tau aggregates uniquely engaged cellular metabolism machinery, including key glycolytic enzymes and TCA cycle proteins, alongside glutamate/GABA neurotransmitter cycling components, with the astrocytic glutamate transporter SLC1A2 showing 27-fold enrichment over other tauopathies. In contrast, PSP tau displayed the most distinctive profile, with extensive protein depletion (52/57 significant proteins) and selective enrichment of proteasome components, particularly PSMB7 showing >3000-fold abundance. DLB tau is associated with neurogenesis modulators while depleting neuroinflammatory mediators. These interaction patterns were validated through proximity ligation assays and correlated with distinct post-translational modification profiles, with PSP tau exhibiting globally elevated ubiquitination, AD showing mixed modification patterns, and DLB displaying minimal ubiquitination. Critically, sarkosyl-soluble fractions revealed reduced interactome complexity across diseases, except for PSP tau which maintained robust interactions with GPCR-ERK signaling and kinetochore proteins, suggesting unique aggregation mechanisms. Our findings establish that conformationally distinct tau strains dictate disease-specific protein interaction networks, providing molecular insight into tauopathy diversity and identifying novel therapeutic targets for precision medicine approaches in neurodegeneration.

## INTRODUCTION

Pathological tau accumulation is strongly linked to neurodegeneration and cognitive decline across neurodegenerative diseases such as Alzheimer’s disease (AD), progressive supranuclear palsy (PSP), dementia with Lewy bodies (DLB), and other tauopathies [10, 19, 67, 75, 92, 99]. Notably, tau aggregates display remarkable structural and functional diversity between tauopathies, with distinct patterns of brain region distribution and cellular targets [13, 24, 33, 38, 63, 86].

PSP is characterized by neuronal and glial tau pathology, mainly involving 4R-tau isoforms forming globose neurofibrillary tangles and tufted astrocytes[20, 41, 63]. In AD, amyloid-β co-pathology synergistically accelerates tau-driven neurodegeneration[5]. DLB shows predominant α-synuclein pathology in limbic and neocortical regions, with tau co-pathology present in over 70% of cases and closely linked to cognitive impairment [16, 36]. Across these disorders, toxic soluble tau aggregates drive disease progression by promoting tau misfolding and aggregation [35, 40, 89].

Recent evidence from our research highlights that tau oligomers isolated from postmortem human brains with AD, DLB, and PSP show distinct disease-specific polymorphs, with differences in structure, morphology, seeding ability, stability, and proteolytic digestion profiles, reflecting differential regulation of neuronal function and gene expression [51].

Beyond their aggregation, protein complexes are key molecular entities assembled into functional modules via protein-protein interactions (PPI) to execute core cellular functions [29, 48]. These complexes prominently regulate essential processes, including DNA replication, transcription, translation, RNA splicing, protein secretion, cell cycle progression, signal transduction, and cellular metabolism[44, 93]. PPIs are particularly crucial within cellular signaling networks; for instance, myriad cellular signals are transmitted through direct interactions between protein substrates and their post-translational modifying enzymes such as kinases/phosphatases, ubiquitinases/deubiquitinases, and acetyltransferases/deacetylases[44, 47, 58, 88]. PPI analysis therefore yields meaningful insights into signal transduction mechanisms.

Recent studies underscore tau’s capacity to interact with varied protein partners, influencing signal transduction and diverse cellular processes[12, 39, 83]. However, it remains unknown if disease-specific tau oligomers possess distinct interactomes. To address this knowledge gap, we conducted a comprehensive investigation of the interactomes of soluble tau aggregates isolated from postmortem human brains with AD, PSP, and DLB.

## RESULTS

### Identification of disease-relevant misfolded tau interactomes

Our previous studies have demonstrated that tau oligomers from PBS- and sarkosyl-soluble fractions significantly differ in post-translational modifications, morphology, and function [51, 68]. Therefore, we investigated the interactome of toxic tau aggregates from the two fractions separately. To do so, tau aggregates and their interacting proteins from the two fractions of AD, PSP, and DLB human brain homogenate (n=3 cases, **Table 1**) were isolated using co-immunoprecipitation (co-IP) with T18 antibody, which is specific for misfolded tau aggregates [50, 53, 66, 68] (**Fig. 1A**). The proteins were then identified and quantified using label-free LC-MS, as described previously [98].

**Figure 1.**
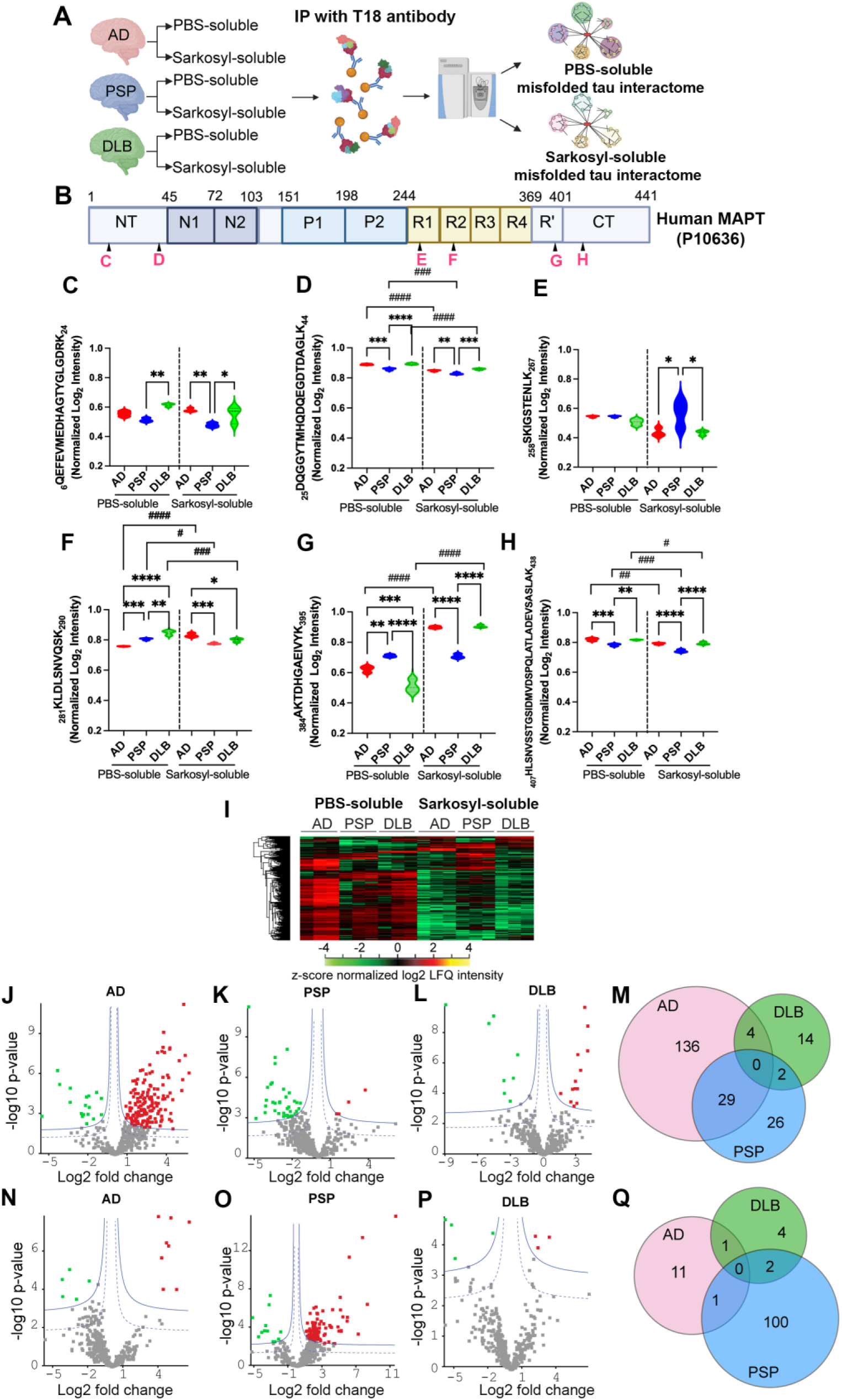
Identification of disease-specific misfolded tau interactomes in PBS- and Sarkosyl-soluble brain fractions from AD, PSP, and DLB. (**A**) Schematic workflow for IP-MS analysis of disease-specific misfolded tau interactome. Misfolded tau was immunoprecipitated from PBS-soluble and Sarkosyl-soluble fractions of human brain (AD, PSP, DLB) using the anti-misfolded-tau antibody T18, followed by label-free liquid chromatography-tandem mass spectrometry (LC-MS/MS) quantification (n = 3). (**B**) Domain map of human MAPT (2N4R; UniProt P10636) with N-terminal region (NT), inserts (N1, N2), proline-rich regions (P1, P2), microtubule-binding repeats (R1–R4 and R′), and C-terminus (CT). Magenta letters mark the positions of representative tryptic peptides plotted in **C–H**. **(C–H**) Normalized log2 LFQ intensities for representative tau peptides detected in misfolded-tau pull- downs, shown for PBS-soluble and Sarkosyl-soluble fractions in AD (red), PSP (blue), and DLB (green). Statistical analyses were calculated by Two-way ANOVA with Tukey test. Asterisks indicate pairwise differences among diseases within a fraction; hash marks indicate PBS- vs Sarkosyl-soluble differences within the same disease (* p<0.05, ** p<0.01, *** p<0.001, **** p<0.0001; # p<0.05, ## p<0.01, ### p<0.001, ### p<0.0001). For (**C**) to (**H**), significant interactors are determined using a permutation-based FDR, and the resulting high-confidence Class A (1% FDR, blue solid line) and low-confidence Class B (5% FDR, blue dashed lines) thresholds are displayed in the plot. Red, proteins significantly enriched (Class A, 1% FDR); green, proteins significantly depleted (Class A, 1% FDR); and grey, non-specific binding proteins. (**I**) Heatmap of significant proteins identified from the pull-down with antibody T18 (ANOVA with permutation-based FDR<5%). **(J–L)** Volcano plots of proteins quantified from PBS-soluble misfolded-tau pull-downs in AD (**J**), PSP (**K**), and DLB (**L**). (**M**) Venn diagram showing PBS-soluble proteins interacting with misfolded tau in each disease. (**N–P**) Volcano plots of proteins quantified from Sarkosyl-soluble misfolded-tau pull-downs in AD (**N**), PSP (**O**), and DLB (**P**). (**Q**) Venn diagram showing Sarkosyl-soluble proteins interacting with misfolded tau in each disease.

**Table 1:**
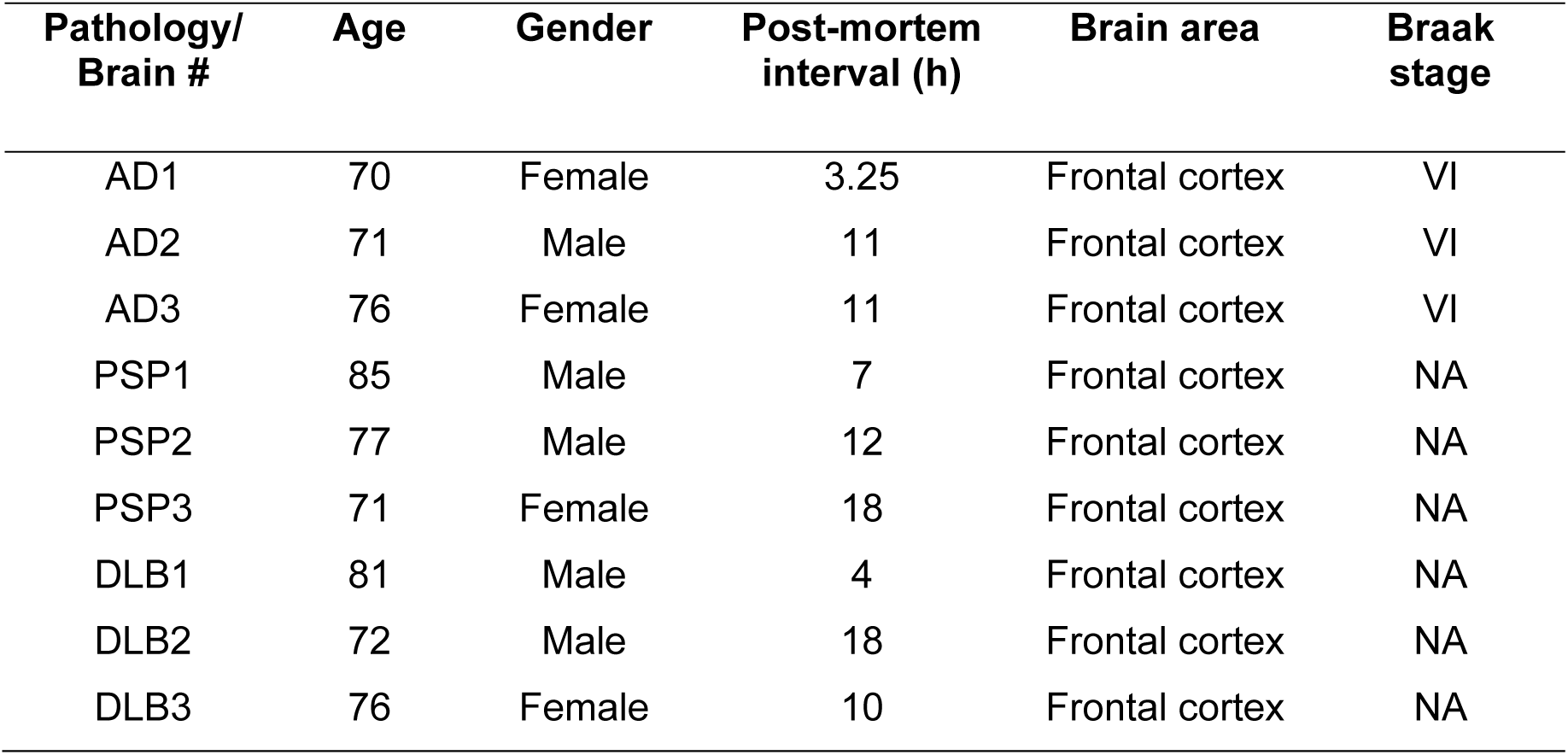
Summary of human brain tissues analyzed in this study.

First, we investigated whether misfolded tau aggregates in AD, PSP, and DLB differ in their susceptibility to trypsin digestion. LC-MS/MS analysis of the tryptic digests of misfolded tau aggregates pulled down from the brains of AD, PSP, and DLB identified 39 tau peptides. To demonstrate the variation in peptide abundance across different tau domains, we normalized the intensity of each peptide with total tau Intensity (**Sup. Fig. 1A**). **Figures 1C-H** show the normalized intensities of peptides mapped to tau domains. Statistical analysis revealed significant differences in intensity of tau peptides between the diseases as well as tau aggregates with different solubility (PBS vs sarkosyl), including the tau fragments from the N-terminal region (NT) (**Fig. 1B-D**), microtube-binding repeats R1 (**Fig. 1B, E**), R2 (**Fig. 1B, F**), R’ (**Fig. 1B, G**), and C-terminal region (**Fig. 1B, H**). Additionally, as illustrated in the MAPT domain mapping (**Sup. Fig. 1B**), we identify seven tau peptides from the PBS-soluble-enriched fraction (**Sup. Fig. 1C-I**) that are mainly located on microtubule-binding regions (R1-R4), and two tau peptides from the sarkosyl-soluble-enriched fraction (**Sup. Fig. 1J-L**) on proline-rich (P2) and R’ regions. These results suggest that misfolded tau aggregates in the brains of AD, PSP, and DBL may have different conformations, leading to variations in trypsin accessibility.

Next, we investigated the disease-specific tau-interactome. Proteomics analysis of tau pull-down identified a total of 493 proteins with high confidence (false discovery rate (FDR) < 1%) **(Sup. Table 1).** Since the abundance of tau varies in each sample, the intensity of each protein was normalized using total tau. Then, multiple ANOVA test with permutation-based FDR was used to identify significant proteins among groups, revealing that 317 proteins exhibited group-wise differences (FDR<5%) (**Fig. 1I**) **(Sup. Table 2)**. The heatmap of these 317 proteins displays the intensity of the proteins in each sample, with proteins in red being enriched in each tau aggregate pull-down and proteins in green being depleted from tau aggregates, compared to other groups. As shown in **Fig. 1I**, significant differences in the misfolded tau interactome are present not only among diseases, but also between the two fractions (PBS vs sarkosyl).

Hawaii plot analysis (multi-volcano plots) with the complement negative control was used to identify the disease-specific misfolded tau interactome [74] (**Fig. 1J-P**). Different s0 and permutation-based FDR parameters were used in the multi-volcano analysis to define class A (higher confidence, s0 = 1, FDR = 1%) and class B (lower confidence, s0 = 1, FDR = 5%) interactors. Each volcano plot displays the results of a pull-down of PBS- or sarkosyl-soluble misfolded tau aggregates from AD (**Fig. 1J, N**), PSP (**Fig. 1K, O**), and DLB (**Fig. 1L, P**), respectively. The thresholds for Class A significant interactors (blue solid lines) and Class B significant interactors (blue dashed lines) are shown in the plot. Class A proteins that were significantly enriched or depleted from PBS-soluble AD derived tau aggregates, relative to PBS-soluble tau aggregates from DLB and PSP, are highlighted in red and green, respectively, while those in grey represent non-specific binding proteins. Only Class A interactors were used for the downstream protein interaction network and bioinformatics analysis. In PBS-soluble fraction, we identified significant Class A interactors of 169 (151 enriched, 18 depleted) from AD (**Fig. 1J**), 57 (5 enriched, 52 depleted) from PSP (**Fig. 1K**), and 20 (12 enriched, 8 depleted) from DLB (**Fig. 1L**), respectively. Similarly, we identified 13 (8 enriched, 5 depleted), 7 (4 enriched, 3 depleted), and 103 (91 enriched, 12 depleted) significant proteins from the AD (**Fig. 1N**), PSP (**Fig. 1O**), and DLB (**Fig. 1P**) pull-downs in the sarkosyl-soluble fraction. After performing Hawaii plot analysis to assess the overall profile of tau interactors, we constructed a Venn diagram to compare the protein sets identified in different sample conditions. Following Hawaii plot analysis for the visualization of the tau interactome profile, Venn diagrams were generated to quantify the number of misfolded tau interactors within the disease-specific interactors, as well as to identify the common interactors among AD, PSP, and DLB from PBS-soluble (**Fig. 1M, Sup. Table 3**) and sarkosyl-soluble fractions (**Fig. 1Q, Sup. Table 4**). Interestingly, there were no common interactors between the three diseases. These results suggest the distinct tau conformation between AD, PSP, and DLB may influence the misfolded tau interacting partners, resulting in diverse pathological phenotypes among the three diseases.

To determine whether the significant disease-specific enrichment or depletion observed in misfolded tau pull-downs was caused by the differential expression levels of these proteins in the brains of patients with AD, PSP, and DLB, we conducted a proteomics analysis of the whole lysate of the brain tissues. We compared the results with those from the misfolded tau pull-down experiments. For example, among the 169 significant proteins identified in the PBS-soluble misfolded AD derived tau pull-down, 119 were also quantified in the proteome study. As shown in **Sup. Fig. 1M**, there is no strong correlation between the disease-specific changes of these proteins in PBS-soluble misfolded tau pull-downs and their differential expression in the brain tissues, similar to the sarkosyl soluble misfolded tau pull-downs (**Sup. Fig. 1N**). Similarly, there is no strong correlation between the PSP or DLB-specific changes of tau interactors and their differential expression in the brain tissues (**Sup. Fig. 1M, N**, **Sup. Table 5)**. Western blot validation of the representative tau interactors related to the TCA cycle (FH) and glucose metabolic process (FABP5) shows that their expression levels were not significantly different in the brains of AD, PSP, and DLB, or between the two fractions. (**Sup. Fig.1O-Q**). Together, these results suggested that the significant changes observed in misfolded tau pull-downs were caused by alterations in protein-protein interactions, rather than differential expression levels in the brains of patients with AD, PSP, and DLB.

### Disease-specific PBS-soluble tau interactomes reveal distinct functional signatures

Following Hawaii plot analysis of PBS-soluble proteins, functional enrichment and network annotation using Cytoscape StringApp [21] and EnrichmentMap tool [3, 39] revealed that the 169 significant proteins are predominantly associated with cellular metabolism, synaptic function, neuron development, and inflammatory response (**Fig. 2A**). Clusters enriched in glucose metabolism, TCA cycle, and respiratory electron transport processes were detected among PBS-soluble AD tau aggregates, with network analysis (**Fig. 2B**) highlighting co-immunoprecipitation of key glycolytic enzymes (HK1, PKM, PFKM) and mitochondrial proteins (PHDX, CS, ACO2, MDH2, FH, ETFA, CYCS) with misfolded AD tau. These findings underscore strong links between AD tau pathology and early metabolic dysfunction [9, 37, 49, 69].

**Figure 2.**
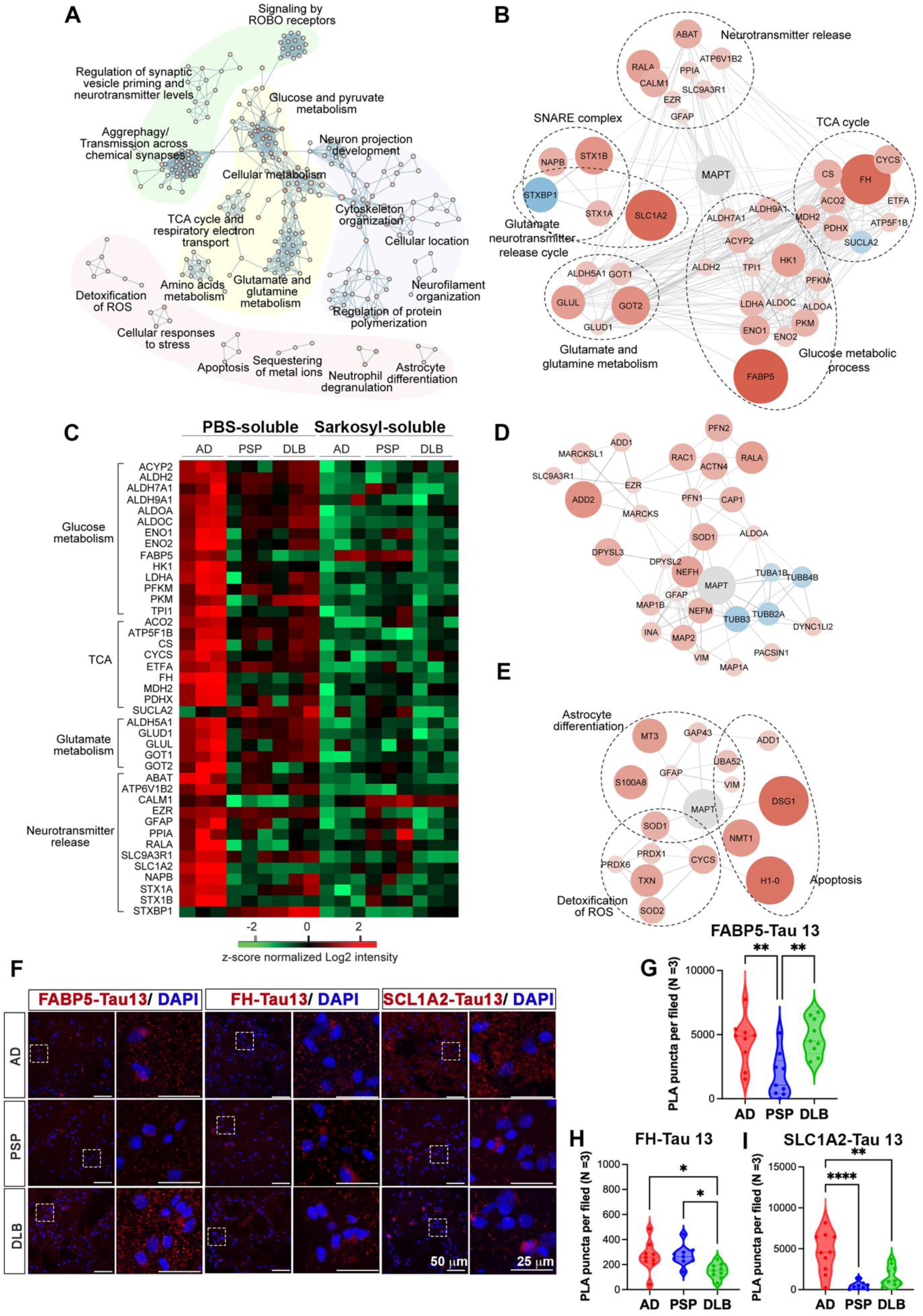
AD misfolded-tau interactome in the PBS-soluble fraction. (**A**) Functional enrichment of the 169 significant proteins either enriched with, or depleted from, misfolded tau immunoprecipitated from PBS-soluble AD brain extracts. (**B**) Protein networks of significant AD misfolded-tau interactors involved in glucose metabolism, the TCA cycle, glutamine and glutamate metabolism and release, the SNARE complex, and neurotransmitter release. Red indicates enrichment with misfolded-tau from AD, while blue indicates depletion. The size of the nodes correlates with the magnitude of the fold changes. (**C**) Heatmap of significant AD misfolded-tau interactors involved in (**B**), shown as z-score–normalized Log2 LFQ intensities across PBS- and sarkosyl-soluble fractions from AD, PSP, and DLB. (**D**) Subnetwork of significant AD misfolded-tau interactors involved in cytoskeleton organization. (**E**) Subnetwork of significant AD misfolded-tau interactors involved in detoxification of ROS, astrocyte differentiation, and apoptosis. (**F**) Representative of *In situ* proximity ligation assays (PLA) using Tau13 (total tau) with FABP5, FH, or SLC1A2, followed by confocal image analysis in AD, PSP, and DLB human brain tissues. Scale bars: 50 μm (main panels), 25 μm (zoom, right panels). (**G-I**) Quantification of PLA puncta per field for (**G**) FABP5–Tau13, (**H**) FH–Tau13, and (**I**) SLC1A2–Tau13 interactions in AD (red), PSP (blue), and DLB (green) (n = 3). Data presented as mean ± SEM. *p < 0.05, **p < 0.01, ****p < 0.0001 by one-way ANOVA with Tukey test.

Interactome analysis further revealed that key enzymes of the glutamate/GABA-glutamine cycle, including GOT1, GOT2, GLUD1, GLUL, and ALDH5A1, as well as SNARE complex proteins (STX1A, STX1B, STXBP1, NAPB) (**Sup. Fig. 1M**) and glutamate transporter SLC1A2 were highly enriched with AD tau aggregates, showing ∼27-fold greater SLC1A2 interaction with PBS-soluble AD tau than DLB or PSP tau (**Fig. 2C**). These interactions imply tau aggregates in glutamate and GABA metabolism and neurotransmitter regulation. Notably, these associations were specific to PBS-soluble AD tau aggregates, with no comparable interactions detected from sarkosyl-soluble AD tau or tau aggregates from PSP or DLB. Additionally, PBS-soluble AD tau showed enhanced interaction with cytoskeletal proteins (GFAP, NEFH, NEFM, MAP1A, MAP1B), while several tubulins (TUBA1B, TUBB2A, TUBB3, TUBB4B) were depleted (**Fig. 2D**). Furthermore, PBS-soluble AD tau was enriched for proteins related to ROS detoxification (TXN, PRDX1, SOD1, SOD2), astrocyte differentiation, and apoptosis (**Fig. 2E**). Validation by proximity ligation assay (PLA) in AD brain tissues confirmed significantly more PLA puncta for FABP5 (metabolism), FH (TCA cycle), and SLC1A2 (glutamate transporter) in AD versus PSP and DLB (**Fig. 2F–I**), with dot blot analysis confirming these interactions primarily with PBS-soluble tau (**Sup. Fig. 2G–J**).

In marked contrast to the metabolically focused AD tau interactome, functional enrichment analysis of DLB tau-associated proteins (20 significant proteins) identified three distinct annotation clusters: astrocyte development, neurogenesis, and vesicle docking (**Fig. 3A**). Notably, neuroinflammatory mediators, S100A8, S100A9, and metallothionein-3 (MT3) were significantly depleted from DLB tau compared to AD and PSP tau, while neurogenesis-related proteins (1N4R tau isoform, GAS7, APOE, BAIAP2) were enriched, and NEFL was depleted (**Fig. 3B**).

**Figure 3.**
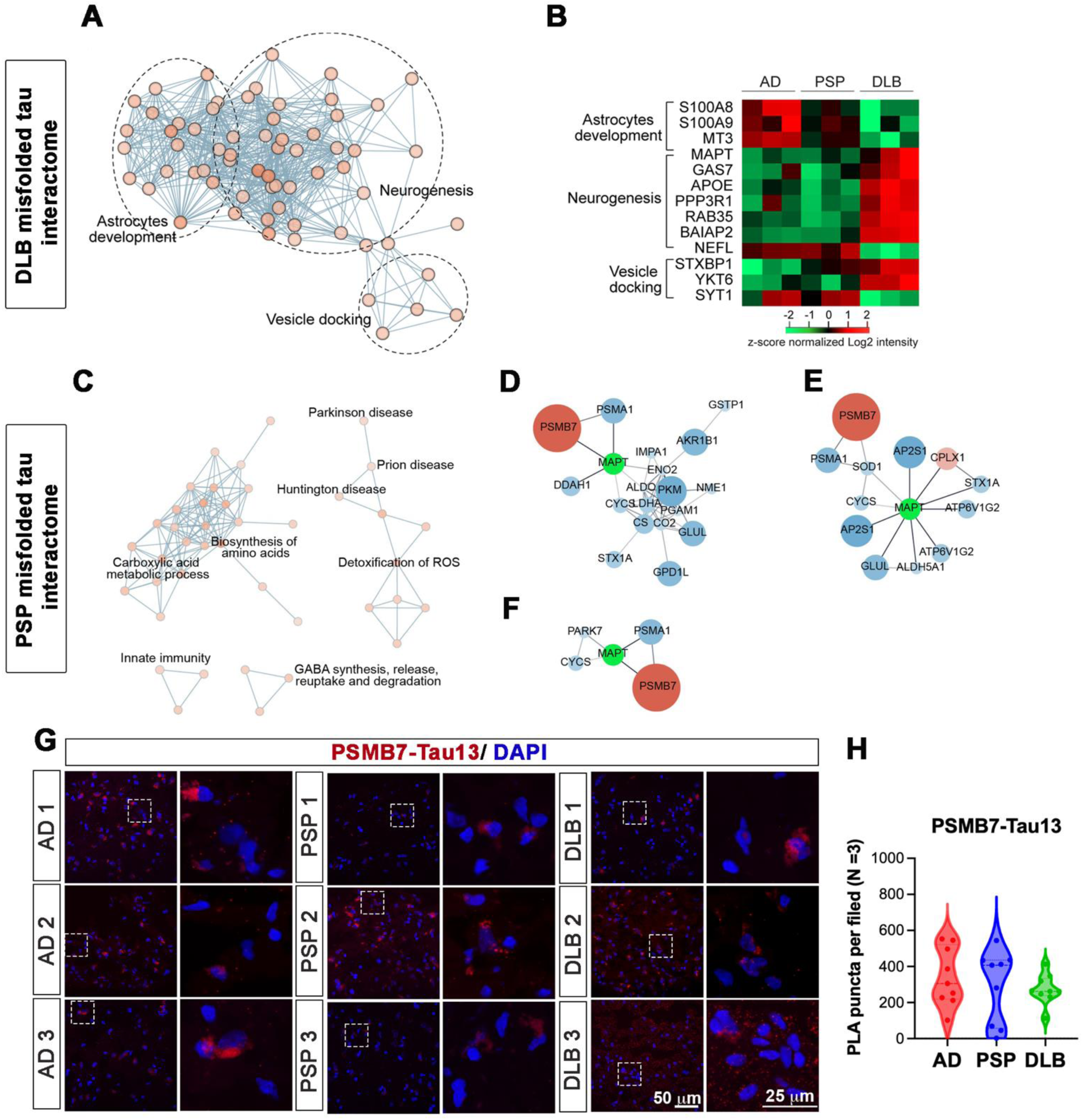
DLB and PSP misfolded-tau interactomes in the PBS-soluble fraction. (**A**) Functional enrichment analysis of the 20 significant proteins either enriched with or depleted from misfolded tau isolated from DLB cases. (**B**) Heatmap of significant DLB misfolded tau interactors involved in astrocyte development, neurogenesis, and vesicle docking. (**C**) Functional enrichment of the 57 significant proteins either enriched with, or depleted from, PSP misfolded tau in PBS-soluble fraction. (**D-F**) Protein networks of significant PSP misfolded tau interactors involved in (**D**) cellular metabolism, (**E**) GABA synthesis, release, reuptake and degradation, as well as synaptic vesicle cycle, and (**F**) Parkinson’s disease. Red indicates enrichment with PSP misfolded-tau, while blue indicates depletion. The size of the nodes correlates with the magnitude of the fold changes. (**G**) PLA images showing colocalization of Tau13 with PSMB7 across AD, PSP, and DLB brain samples. Scale bars: 50 μm (left panels), 25 μm (zoom, right panels). (**H**) Quantification of PLA puncta per field for PSMB7–Tau13 across disease groups (n = 3). Data shown as mean ± SEM, analyzed by one- ANOVA with Tukey test.

PSP tau exhibited the most distinctive interactome profile, with only 5 of 57 significant proteins showing enrichment (TRIM21, PSMB7, TBC1D10B), while the majority (52/57) were depleted compared to AD and DLB tau. These proteins were functionally associated with cellular metabolism, stress response/ROS detoxification, GABA neurotransmission, and innate immunity (**Fig. 3C**). In direct opposition to AD tau findings, metabolic proteins were predominantly depleted from PSP tau (**Fig. 3D**), as were proteins involved in GABA synthesis, release, and degradation, except for PSMB7, verified by PLA assay (**Fig. 3G-H**), and CLPX1 (**Fig. 3E**). Several Parkinson’s disease-associated proteins, including PARK7, were also depleted from PSP tau (**Fig. 3F**).

### Sarkosyl-soluble tau fractions exhibit reduced interactome complexity with disease-specific signatures

While PBS-soluble tau fractions demonstrated extensive protein interactions, analysis of the sarkosyl- soluble fractions revealed markedly fewer significant protein associations across all three diseases. MS analysis identified only a limited number of proteins significantly enriched or depleted from sarkosyl- soluble AD or DLB tau, with corresponding protein networks shown in **Fig. 4A, B**. Although functional enrichment analysis of these proteins did not yield significantly enriched terms, functionally important proteins were identified, including ubiquitin carboxyl-terminal hydrolase isozyme L1 (UCHL1), which was depleted from sarkosyl-soluble AD tau, and E3 ubiquitin-protein ligase MARCH9, which was depleted from sarkosyl-soluble DLB tau.

**Figure 4.**
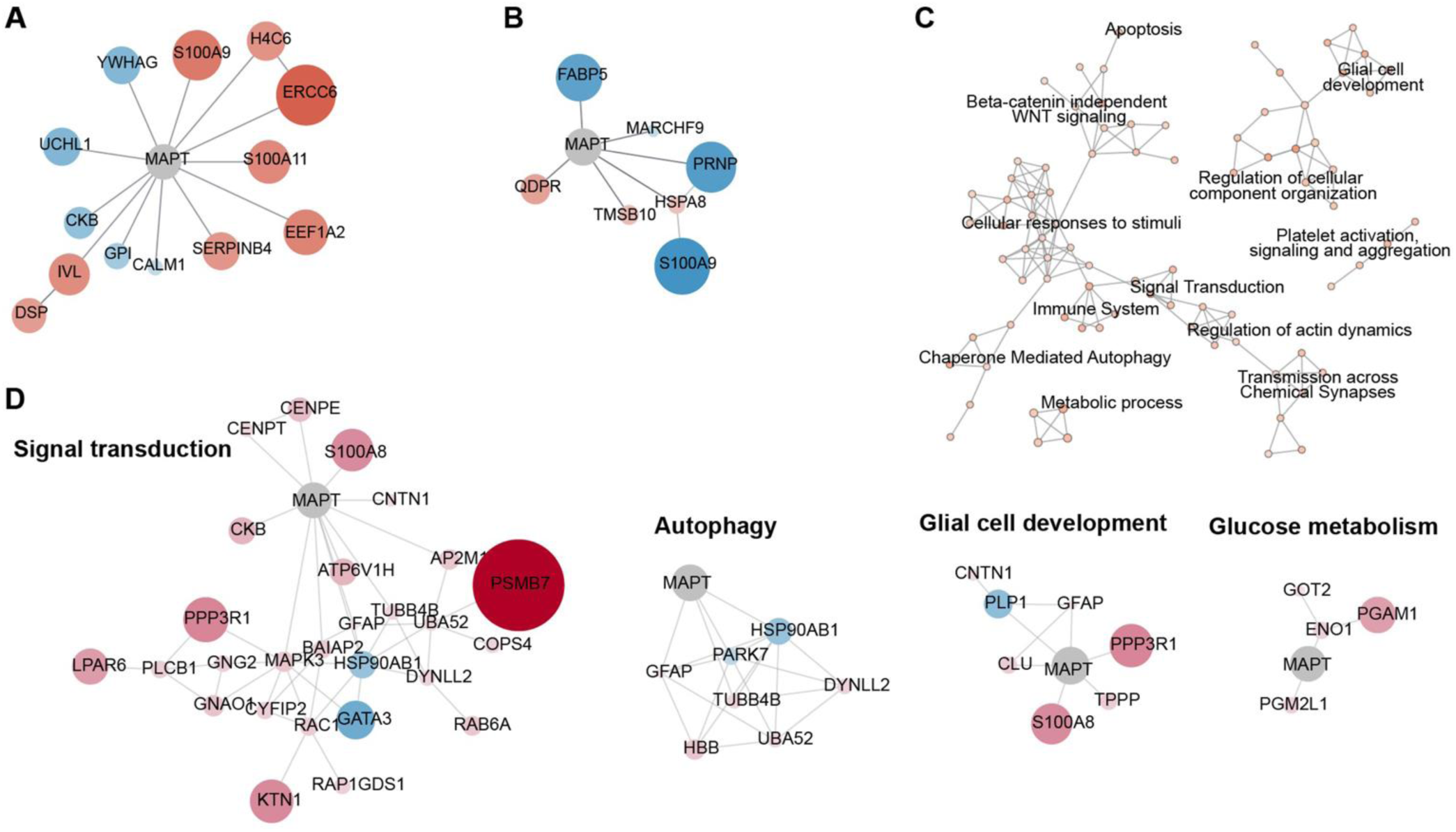
AD, DLB, and PSP misfolded tau interactomes in sarkosyl-soluble fraction. (**A**) Protein networks of sarkosyl-soluble AD misfolded-tau interactors. Red indicates enrichment with AD misfolded tau, while blue indicates depletion. The size of the nodes correlates with the magnitude of the fold changes. (**B**) Protein networks of sarkosyl-soluble DLB misfolded-tau interactors. (**C**) Functional enrichment analysis of the 103 significant proteins either enriched with sarkosyl-soluble PSP misfolded-tau or depleted from it. (**D**) Protein networks of significant PSP misfolded-tau interactors involved signaling transduction, autophagy, glial cell development, and glucose metabolism.

In contrast to AD and DLB, sarkosyl-soluble PSP tau demonstrated a more robust interactome, with MS analysis identifying 103 significant proteins that yielded multiple significantly enriched annotation terms (**Fig. 4C**). Notably, sarkosyl-soluble PSP tau interacted with proteins involved in GPCR-ERK pathway signaling (MAPK3, GNG2, GNAO, PLCB1, PPR3R1, and LPAR6) and kinetochore function (CENPT and CENPE) (**Fig. 4D**). Additional interactions were observed with proteins regulating autophagy, glial cell development, and cellular metabolism (**Fig. 4D**), suggesting that PSP tau maintains distinct functional associations even in the more in soluble sarkosyl-extracted fraction.

### Machine learning classification identifies disease-specific tau interactome signatures

Machine learning analysis was employed to identify the most discriminative features of tau interactomes across disease phenotypes. In the PBS-soluble fraction, ANOVA identified 264 significantly different proteins (permutation-based FDR <0.01), and Support Vector Machine classification with 4-fold cross- validation achieved 0% prediction error using only the top 4 features (**Fig. 5A**). Principal component analysis of the top 10 proteins (**Table 2, Sup. Fig. 3, Fig. 5B**) demonstrated distinct segregation of AD, PSP, and DLB cases (**Fig. 5C**), with PCA loadings revealing disease-specific contributions (**Fig. 5D**). SLC1A2 and STX1B interactions were most distinctive for AD (38× and 13× more abundant than DLB/PSP, respectively), BAIAP2 and RAB35 characterized DLB, while PSMB7 and MARS distinguished PSP.

**Figure 5.**
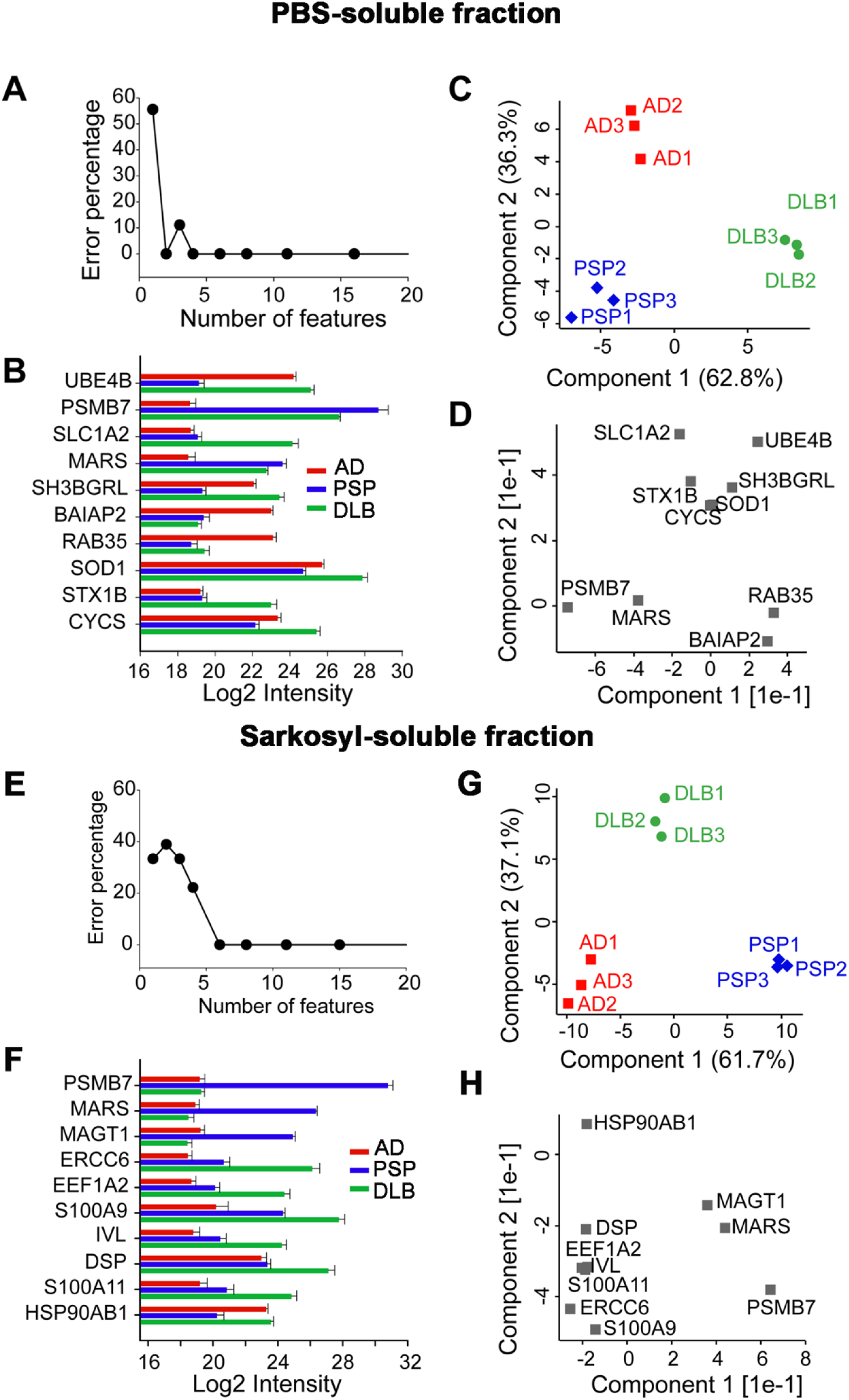
Classification analysis of disease-relevant misfolded-tau interactomes in PBS- and sarkosyl-soluble fraction. (**A, E**) The prediction error percentage corresponding to the number of features used for disease phenotype prediction in (**A**) PBS-soluble or (**E**) sarkosyl-soluble fraction by using the SVM algorithm. (**B, F**) Abundance (Log2 LFQ intensity) of the ten most informative proteins in (**B**) PBS-soluble or (**F**) sarkosyl-soluble fraction across AD (red), PSP (blue), and DLB (green) misfolded-tau interactomes. (**C, G**) PCA analysis using the abundances of these top ten proteins in (**C**) PBS-soluble or (**G**) sarkosyl- soluble fraction across AD (red), PSP (blue), and DLB (green) misfolded-tau interactomes. (**D, H**) The PCA loadings of the top ten proteins in (**D**) PBS-soluble or (**H**) sarkosyl-soluble fraction. It illustrates how much each protein influences the variance captured by that component.

**Table 2:**
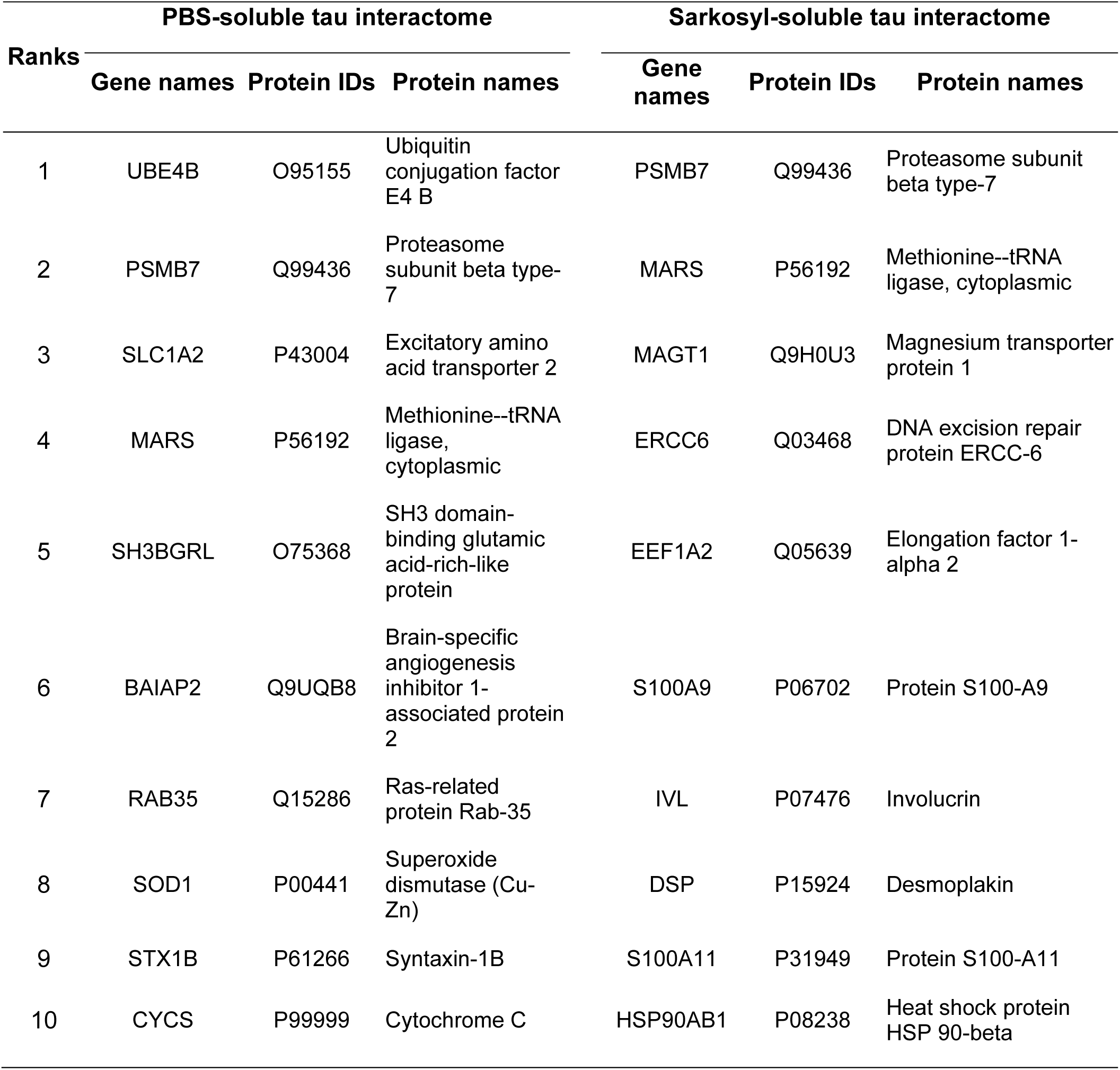
Top 10 features of disease-relevant misfolded tau interactors in PBS-soluble and Sarkosyl-soluble fractions.

Similarly, in the sarkosyl-soluble fraction, 166 significant proteins were identified, with the top 6 features achieving 0% classification error (**Fig. 5E**). PCA of the top 10 proteins (**Table 2, Sup. Fig. 3, Fig. 5F**) again showed clear disease separation (**Fig. 5G**). Notably, PSMB7 and MARS remained highly distinctive for PSP across both fractions, with PSMB7 showing >3000-fold enrichment in sarkosyl-soluble PSP tau compared to other diseases (**Fig. 5H**). AD was characterized by interactions with ERCC6, EEF1A2, and S100A11, while HSP90AB1 was strongly associated with DLB. These findings demonstrate that tau interactomes contain robust, disease-specific molecular signatures that enable accurate phenotypic classification.

### Ubiquitination enzymes modify disease-specific tau conformations

Proteomic profiling identified 20 ubiquitinated tau peptides across AD, PSP, and DLB, with 15 peptides exhibiting significant disease-specific differences after normalization to total tau (**Fig. 6A, Sup. Fig. 5**). In PBS-soluble fractions, ubiquitination at Lys-24, Lys-87, Lys-267, Lys-300, Lys-311, Lys-338, and Lys-385 (2N4R/2N3R isoforms) was elevated in AD tau compared to DLB but remained lower than in PSP tau. PSP tau showed globally elevated ubiquitination in both PBS- and sarkosyl-soluble fractions, while DLB tau displayed the lowest levels except for Lys-254 (2N4R), which was specifically increased. In sarkosyl-soluble fractions, most AD tau ubiquitination resembled DLB, except for Lys-254, which was higher in AD than PSP.

**Figure 6.**
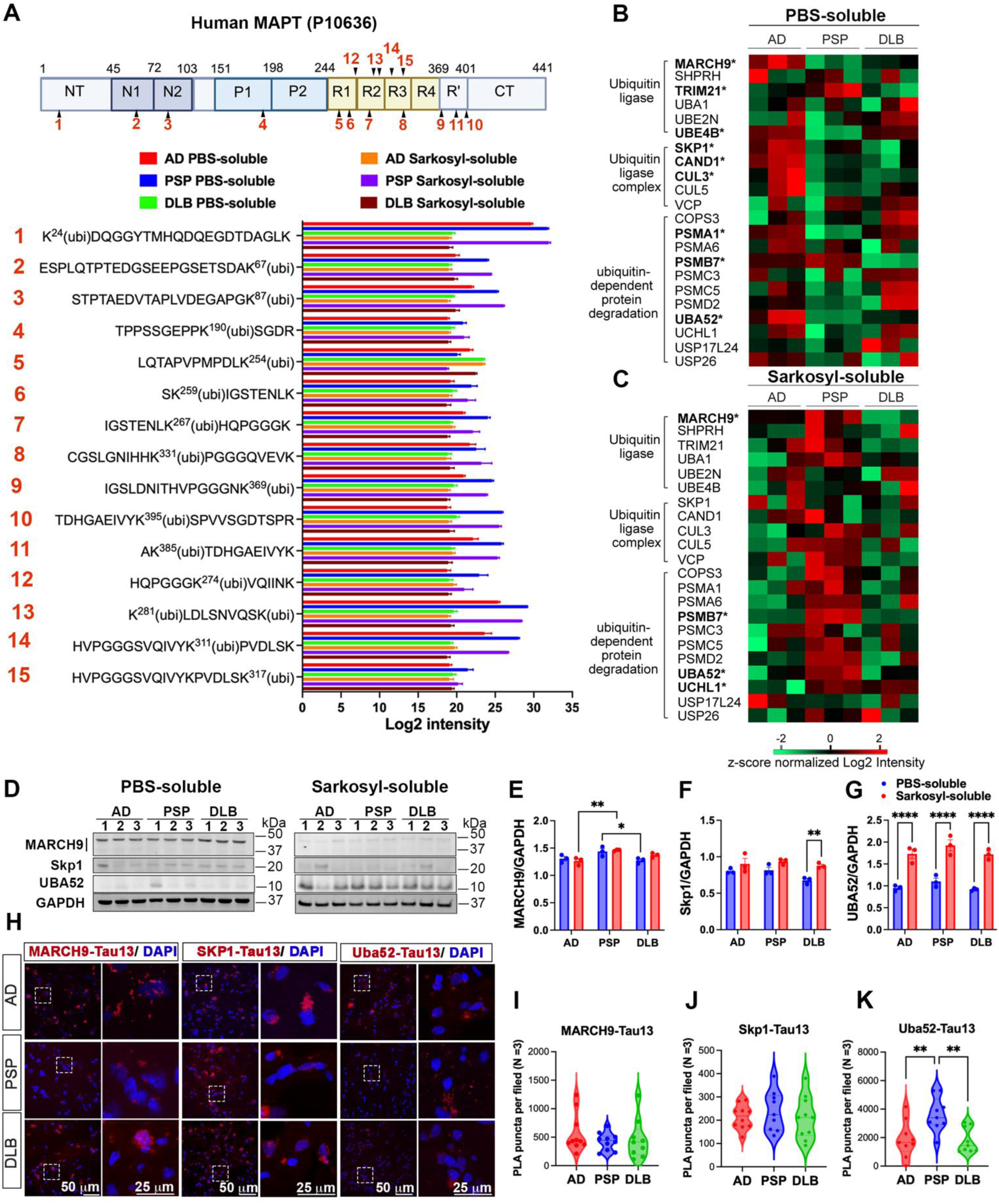
Ubiquitination of misfolded tau and ubiquitination-related interactors. (**A**) Domain map of human MAPT (UniProt P10636) showing its ubiquitinated lysine residues (1-15 in red) mapped above the schematic (top) and its respective quantification (bottom). Profiles of 15 ubiquitinated tau peptides that exhibited significant variance in a disease-specific manner (ANOVA with permutation- based FDR<0.01). (**B–C**) Heatmaps of ubiquitination-related interactors of misfolded-tau from (**B**) PBS-soluble and (**C**) sarkosyl-soluble fractions, grouped as ubiquitin ligases, ubiquitin ligase complex, and ubiquitin- dependent protein degradation. Proteins shown in bold and marked with * are significant by ANOVA with permutation-based FDR < 0.01 (z-score–normalized log₂ intensity across AD, PSP, and DLB). (**D-G**) Representative immunoblots and densitometric quantification of (**E**) MARCH9 (ubiquitin E3 ligase), (**F**) SKP1 (SCF complex), and (**G**) UBA52 (ubiquitin–ribosomal fusion protein) normalized to GAPDH in PBS- and Sarkosyl-soluble brain fractions (AD, PSP, DLB; n = 3 per group). Quantification of band intensity was normalized to GAPDH. The same immunoblots probed with loading controls shown in **Sup. Fig. 1** and **3**. Data are mean ± SEM with individual cases overlaid. *p < 0.05, **p < 0.01, ****p < 0.0001 by two-way ANOVA with Tukey Test (**H-K**) Representative PLA images and quantification of PLA puncta per field showing colocalization of Tau13 with (**I**) MARCH9, (**J**) Skp1, and (**K**) UBA52 in postmortem brain tissue from AD, PSP, and DLB. Boxed regions are magnified at right. Scale bars: 50 μm (left panels), 25 μm (zoom, right panels). Data presented as mean ± SEM. **p < 0.01 by one-way ANOVA with Tukey Test.

Interactome studies revealed that E1 (UBA1) and E2 (UBE2N) ubiquitin enzymes did not show disease-specific interactions, whereas E3 ligases were differentially associated. In the PBS-soluble fraction, AD tau specifically interacted with E3 ligases MARCH9 and UBE4B, while PSP tau was enriched for TRIM21. DLB tau lacked significant E3 interaction, consistent with its low ubiquitination profile. In the sarkosyl-soluble fraction, PSP tau uniquely enriched for MARCH9; AD and DLB tau did not display significant E3 associations (**Fig. 6B, C**). PBS-soluble AD tau also associated with cullin-RING ligase complex components CUL3, SKP1, and CAND1, implicating the BCR and SCF complexes in AD-specific ubiquitination. Several key substrates of these complexes, including NFKBIB, NFKBIE, and others involved in signaling and cell cycle, were identified as potential targets.

Deubiquitinating enzymes UCHL1, USP17L24, and USP26 did not show disease-specific interactions with tau aggregates in the PBS-soluble fraction; however, in the sarkosyl-soluble fraction, UCHL1 binding to AD tau was significantly decreased compared to DLB and PSP, which may influence the elevated ubiquitination at Lys-254 in insoluble AD tau.

The tau interactome also showed that only PSP tau aggregates, in both PBS- and sarkosyl-soluble fractions, interacted significantly with PSMB7 (20S proteasome core), while AD and DLB tau did not exhibit substantial proteasome association (**Fig. 6B, C**). This unique association suggests differential proteasomal targeting among tauopathies. Confirmation by western blot, PLA, and dot blot (**Fig. 6D–K, Sup. Fig. 4**) corroborated the interactome findings regarding E3 ligases, SCF components, and proteasome subunits.

Collectively, these data demonstrate that tau aggregates from AD, PSP, and DLB display distinct profiles of ubiquitination, shaped primarily by differential E3 ligase interactions and supported by unique associations with ligase complexes and the proteasome system.

### Distinct acetylation and phosphorylation profiles define disease-specific tau aggregates

Tau lysine residues are subject to acetylation, which can compete with ubiquitination at the same sites and influence protein stability. Proteomic analysis identified seven acetylation sites on tau, with Lys-24, Lys-163, Lys-267, and Lys-281 showing significant disease-specific differences (**Fig. 7A–E**). Acetylation of Lys-267, Lys-281, and Lys-163 was notably lower in PSP tau aggregates compared to AD and DLB, whereas acetylation of Lys-24 was significantly elevated in PBS-soluble AD and PSP tau, as well as in sarkosyl-soluble PSP tau. These profiles were validated by PRM-MS (**Sup. Fig. 5**). No specific acetyltransferase or deacetylase was found to be associated with tau aggregates in the interactome analysis, leaving the enzymatic basis of these modifications unresolved.

**Figure 7.**
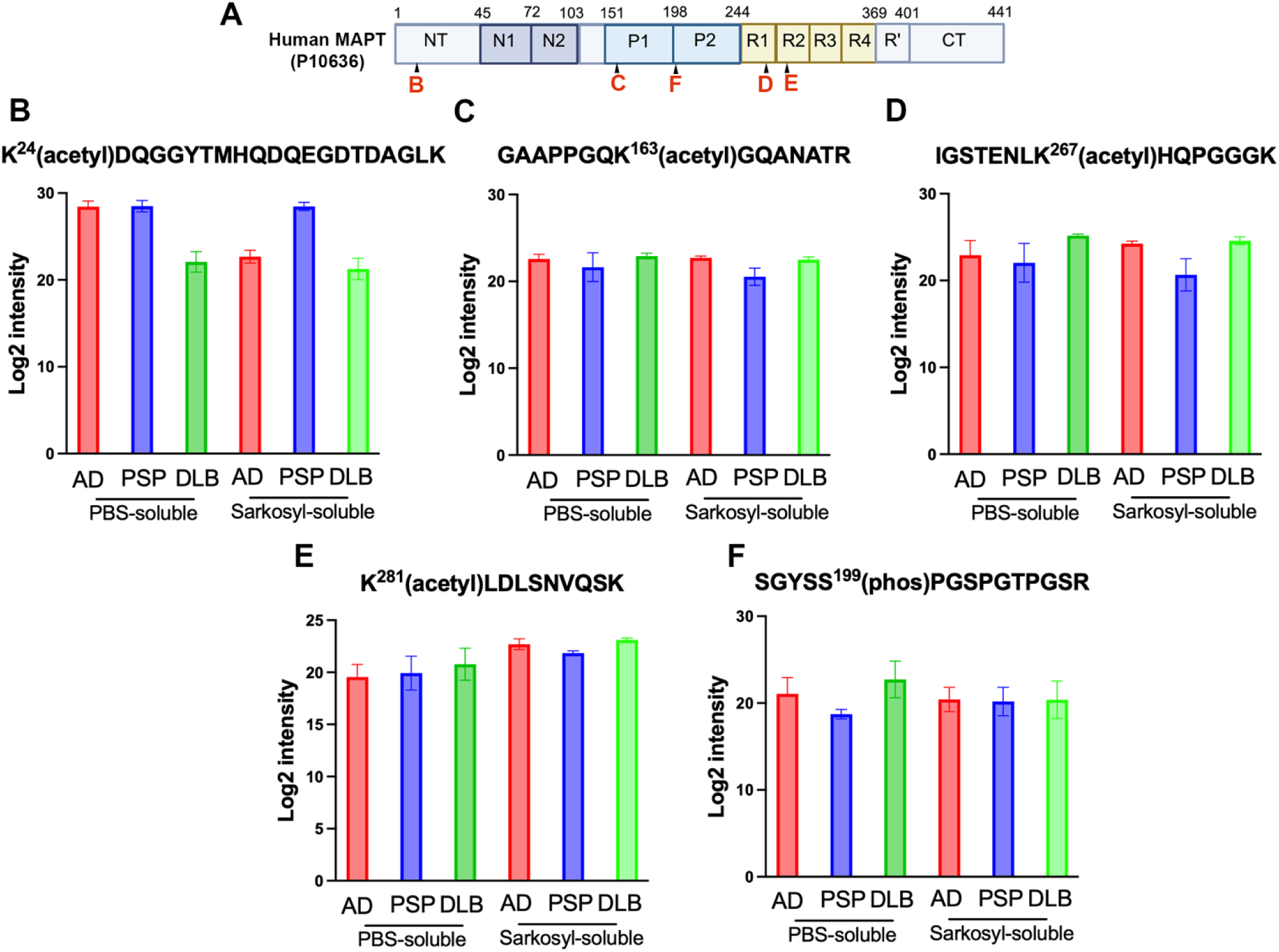
Acetylation and phosphorylation of misfolded tau. (**A**) MAPT domain schematic of the acetylation (**B-E**) and phosphorylation sites (**F**) with its respective quantified peptides that exhibited significant variance in a disease-specific manner (ANOVA with permutation-based FDR<0.01).

In parallel, six phosphorylation sites were detected, with phosphorylation at Ser-199 being markedly higher in AD and DLB tau than in PSP tau (**Fig. 7A, F**), highlighting a disease-specific phosphorylation signature relevant to neurofibrillary degeneration[55, 56].

Together, these findings demonstrate that disease-specific tau polymorphs exhibit distinct patterns of ubiquitination, acetylation, and phosphorylation. Among these modifications, ubiquitination displays the most pronounced disease-specific variation.

## DISCUSSION

This study provides the first comparative, proteomic-based characterization of misfolded tau interactomes isolated from PBS and sarkosyl-soluble fractions of AD, PSP, and DLB post-mortem brain. By coupling conformation-specific capture of misfolded tau (T18) with label-free LC-MS, we uncover three principal and interrelated findings (**Fig. 8**): 1) Tau aggregates form disease-specific interaction networks with distinct molecular and cellular functions; 2) PBS-soluble misfolded AD derived tau harbor richer and more functionally coherent interactomes than sarkosyl-soluble fractions, while sarkosyl-soluble misfolded PSP derived tau forms more protein complexes than PBS-soluble misfolded PSP derived tau. Neither PBS- soluble nor sarkosyl-soluble misfolded DLB derived tau has many interactors; 3) The pattern of PTMs, especially ubiquitination, but also acetylation and phosphorylation, aligns with disease-specific E3 ligase and proteasome associations. Together, these findings underscore that the conformational heterogeneity of tau polymorphs, along with the formation of different tau-protein complexes, drives divergent cellular pathobiology and explains the heterogeneity of tau pathologies.

**Figure 8.**
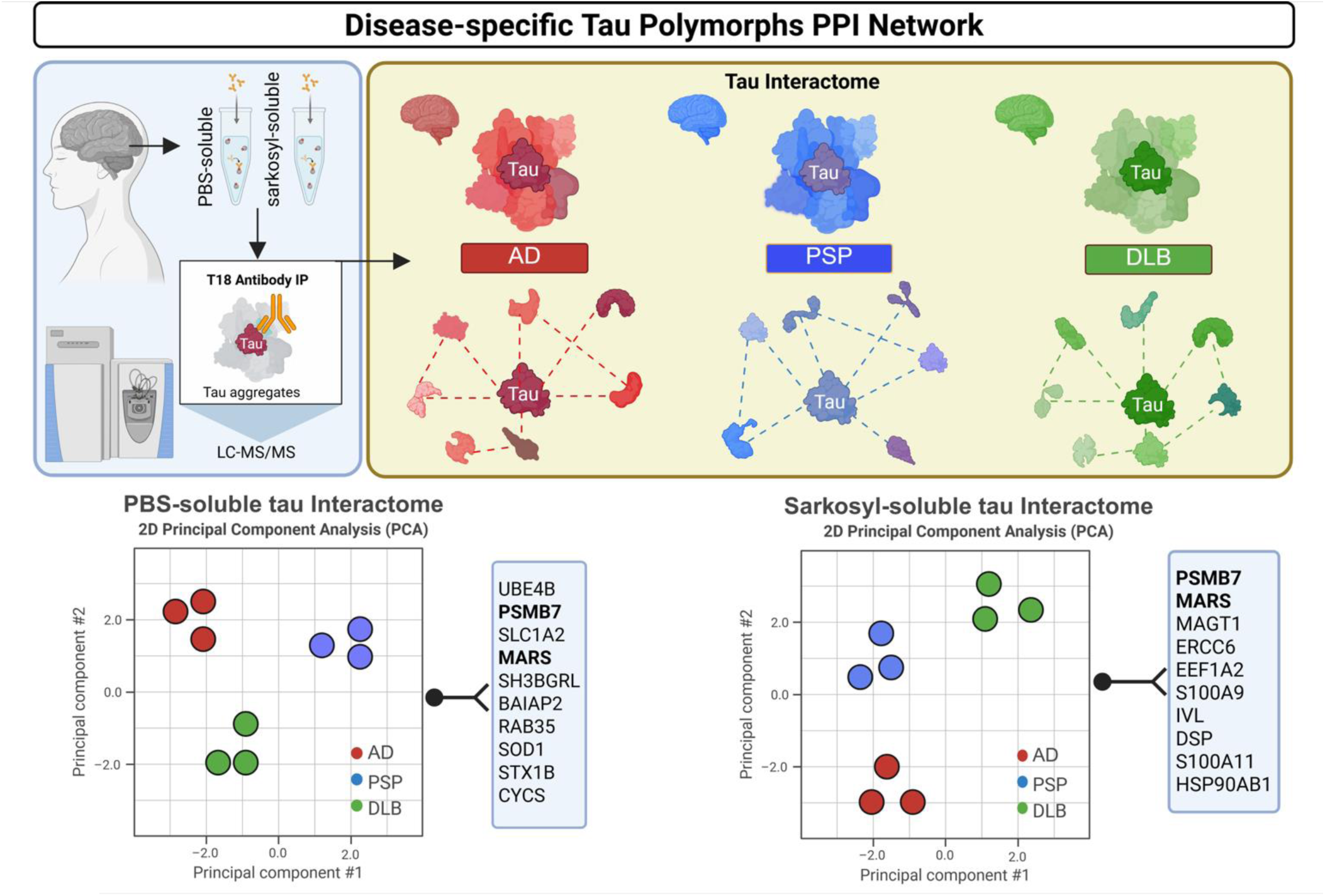
Schematic overview of the study.

The distinct tau interactomes and PTM profiles in the brains of AD, PSP, and DLB support the concept that tau adopts disease-specific conformations that expose different interaction surfaces and therefore recruit different sets of cellular proteins. Our previous as well as high-resolution CyroEM studies have demonstrated disease-specific soluble and filament folds (PHF/SF in AD vs unique folds in several 4R tauopathies), providing compelling evidence for selective interactor recruitment [26, 51, 76]. Moreover, the apparent differences in the PBS soluble fraction suggest that early conformers rather than late are the principal determinants of selective recruitment and early cellular dysfunction [2, 15, 40, 42, 43, 79, 80].

PBS-soluble AD tau selectively co-immunoprecipitated glycolytic enzymes (HK1, PKM, PFKM) and TCA/mitochondrial proteins (CS, ACO2, MDH2, FH, ETFA, CYCS), and strongly enriched glutamate/GABA cycle enzymes and the astrocytic glutamate transporter SLC1A2/EAAT2. This pattern, combined with extensive literature implicating early metabolic and mitochondrial dysfunction in AD, suggests that tau can disrupt mitochondrial dynamics and bioenergetics [9, 32, 70, 78, 83]. The SCLA2/EAAT2 enrichment is particularly striking given prior evidence that EAAT2 dysfunction reduces glutamate clearance in AD and contributes to excitotoxic stress, which is known to be association between soluble tau aggregates, and glutamate-handling machinery provides a mechanistic route linking tau conformers to synaptic hyperexcitability and metabolic stress [9, 83].

PBS soluble tau from AD is also associated with the SNARE complex proteins and syntaxins, implicating misfolded tau in presynaptic vesicle mechanism. Recent interactomes and functional studies map tau binding to presynaptic vesicle proteins and show syntaxins which can facilitate tau secretions, known mechanisms that may underlie trans-synaptic spread and synaptic dysfunction in tauopathies [83]. As such, our data complements other studies showing that misfolded tau may impair energy supply, alter neurotransmitter handling and presynaptic release, producing a convergent synaptic producing a convergent synaptic/metabolic vulnerability in AD.

Enrichment of antioxidant enzymes (TXN, PRDX1, SOD1/SOD2) and astrocytic markers with tau derived from AD supports the model which suggests that misfolded tau may perturb neuron-glia metabolic coupling and antioxidant defenses, a phenomenon also seen with Aβ [65]. Oxidative stress is well-documented in AD and promotes tau pathology and arises as a downstream consequence of mitochondrial dysfunction; the detected direct interaction suggests that misfolded tau may amplify oxidative damage with its association with redox machinery [23, 34].

Comparatively, PSP-derived misfolded tau displayed a strikingly different profile where metabolic and many neurotransmission-associated proteins were depleted from PSP-derived tau (both fractions), while proteostasis-related proteins, particularly PSMB7 and the E3 TRIME21-selectively associated compared to AD or DLB. Consistent with this, PSP phosphorylated tau proximity proteomics independently highlighted enrichment of protein-degradation and stress-response networks alongside cytoskeletal interactors [71]. Of note, PLA confirmed direct interaction with PSMB7 and tau. Although group-wise differences were not statistically significant, the direction and effect sizes were consistent with the interactome data generated through LC-MS/MS. One possible explanation for this is that our LC-MS/MS derives from T18 immunocapture of PBS- and Sarkosyl-soluble fractions, whereas PLA was performed on fixed tissue sections. Fractionation followed by immunocapture enriches misfolded tau containing complexes and boosts signal to noise ratio, while PLA identifies the full cellular milieu, diluting events confined to specific fractions. TRIM21 has been shown to neutralize cytosolic tau seeds by recruiting proteasomal and p97/VCP activity, offering a possible mechanistic interpretation for its PSP association in our data set [52, 62, 71]. This proteostatis-dominant interactome distinguishes PSP from the metabolic/synaptic signature of AD.

DLB associated misfolded tau interactors clustered around vesicle docking, synaptic organization and proteins implicated in neuronal remodeling/neurogenesis (e.g., BAIAP2, GAS7, APOE). The depletion of inflammatory mediators (S100A8/A9), MT3 from DLB derived misfolded tau, contrasting with AD. This pattern fits a broader observation where S100A8/A9 positive microglia are enriched in primary tauopathies and AD as well accelerated innate immune “aging”, while postmortem and in vivo data suggest that robust microglial activation in DLB is often tied to concomitant AD pathology rather than ‘”pure” DLB [1, 31, 90]. MT3, frequently reduced in AD and central to neuronal metal homeostasis, has also been linked to α-syn biology via metalothionien-metal interactions, offering a possible explanation for disease-specific differences in MT3 engagement with tau [7, 59, 84, 96, 100]. Given the frequent copresence and interaction between tau and α-synuclein pathologies, the synaptic/vesicular tilt of the DLB derived tau interactome may capture pathways by which pathologies disrupt presynaptic trafficking and neuronal connectivity. One possible explanation may be that α-syn is concentrated at the presynaptic terminals and pathological increases perturb synaptic vesicular cycling, which is elevated in DLB cohorts [4, 6, 28, 54, 87, 91]. This further explains our tau-centric synaptic/vascular network.

Ubiquitination emerged as the most disease specific as PSP derived tau displayed globally elevated ubiquitination and selective enrichment for proteasomal components, notably TRIM21. This is consistent when previously published works investigating PSP-phospho-tau proximity proteomics showing strong representation of ubiquitin proteosome pathways and with mechanistic works established TRIM21 can neutralize intracellular seeds [57, 62, 71]. By contrast, AD derived tau preferentially associated with cullin- RING ligase components (SKP1, CUL3, CAND1), which aligns with previously published reports of cullin- RING ligase components are activated by neddylation and that neddylation/CRL signaling is perturbed in AD brain [8, 97]. We also report enrichment of ubiquitin E3/E4 factors with AD-derived tau, including MARCH9 and UBE4B. MARCH9 functions as a RING-CH E3 which ubiquitinates membrane proteins to route them to the lysosomes. To our knowledge, direct tau ubiquitination by MARCH9 has not been reported, as such its enrichment in our data extends prior MARCH family involvement beyond MARCH7, which has been reported to ubiquitinate tau and reduce microtubule binding [27]. Interestingly, UBE4B cooperates with CHIP to extend the tau ubiquitin chains and promote clearance [77], providing a potential explanation for their enrichment in AD. Conversely, we also observed the lowest overall tau ubiquitination and minimal E3 enrichment relative AD/PSP.

Mechanistically, our data fits a PTM-cross talk model in which disease specific ubiquitin E3 sculpt tau’s lysine usage and routing. Tau acetylation at key microtubule-binding Lysines play a crucial role in microtubule binding, promoting aggregation, and functionally competes with lysine ubiquitination [14, 25, 60, 94]. Conversely, UBE4B ubiquitination has been reported to suppress aggregation and favors degradation [77]. This fundamentally suggests that E3 interaction and PTM patterning are integral to tau strain identity, aggregation and life span of the aggregates (proteosome vs autophagy) [46, 94]. To add, we did not detect phosphorylation at the AT8 epitope (pSer202/pThr205); however, pSer199 was ∼8-fold more abundant than pSer202 in our dataset. The absence of AT8-site phosphorylation, together with the relative enrichment of pSer199, suggests that the misfolded tau isolated from AD, PSP, and DLB reflects an early pathological stage: pSer199 has been reported as an early marker of tau pathology and is detectable in pretangle neurons, preceding robust AT8 positivity [11, 55, 56, 64].

In summary, our study reveals that misfolded tau is not a single unit rather tau conformers in AD, PSP, and DLB interact with distinct protein networks and PTM states that likely determine cellular pathways disrupted in each disease. These interactomes provide mechanistic hypotheses and nominate testable, disease-specific targets for biomarker development and therapeutic intervention.

## MATERIALS AND METHODS

### Brain homogenate preparation

Postmortem brain tissue samples from AD, PSP, and DLB subjects were obtained from the University of Kentucky Alzheimer’s Disease Center Tissue Bank, Oregon Brain Bank, Michigan Brain Bank, and the Brain Resource Center at Johns Hopkins and approved by the Institutional Ethics Committee. Neuropathological assessment conformed to National Institute on Aging/Reagan Institute consensus criteria. The following information was available for the cases used in this study: diagnosis, age at death, gender, post-mortem index, brain area, and Braak stage (**Table 1**).

### Separation of soluble and insoluble tau

Separation of PBS soluble and insoluble tau was performed in accordance with protocols previously described [68]. In brief, brain tissues and cell lysates were homogenized in 1X phosphate-buffered saline (PBS) with protease inhibitor (Roche) at 1:3 (w/v) ratio and incubated for 20 min at 4 °C. Then the samples were centrifuged at 11,000*g* for 20 min at 4 °C. The supernatant was collected and centrifuged at 100,000*g* for 60 min at 4 °C. To extract the PBS insoluble tau, the pellets from the first and second cold centrifugation were combined and resuspended in PHF extraction buffer (10 mM Tris-HCl.1 mM EGTA, 0.85 M NaCl, 10% (w/v) sucrose, pH 7.4) at 1:10 (w/v) ratio. Samples were centrifuged at 15,000*g* for 20 min at 4 °C. Sarkosyl solution was added to the supernatant at a final concentration of 1% and stirred for 1 h at room temperature (RT) before centrifugation at 100,000*g* for 30 min at 4 °C. Finally, the Sarkosyl insoluble fraction was made by resuspending the pellet in 1X <u>PBS</u> depending on the amount of starting material (1 ml buffer for 25 g of starting material). The total protein concentrations of the final fractions were analyzed by bicinchoninic acid (BCA) assay (Thermo Scientific).

### Immunoprecipitation (IP)

PBS-soluble and Sarkosyl-soluble tau were immunoprecipitated with a misfolded tau specific T18 antibody [50, 53, 66, 68] using Pierce Co-Immunoprecipitation Kit (Thermo Scientific). Briefly, amine- reactive resin was coupled with affinity-purified T18 antibody, incubated with samples. Bound proteins were eluted in 0.1 M glycine (pH 2.8), the pH was adjusted to 7.0 by adding 1 M Tris-HCl (pH 8). Isolated fractions were subjected to buffer exchange against 1X PBS followed by LC-MS/MS and dot blot analysis. The total protein concentration was measured with a BCA assay.

### *In situ* proximity ligation assay (PLA)

To identify the interaction of tau and its interactors in postmortem human brain tissues, the Duolink *In Situ* Red starter kit mouse/rabbit (Sigma Aldrich; DUO92101) was used according to the manufacturer’s guideline. Briefly, frozen human brain tissues with AD, PSP, and DLB were pretreated with chilled methanol before incubating with blocking solution at 37 °C. After 1 h, primary antibody solution was added overnight at 4 °C. Primary antibodies used include mouse anti-Tau 13 (1:20000, BioLegend; MMS-520R) and rabbit anti-interactors, including anti-FABP5 (1:2000, ProteinTech, 12348-1-1AP), anti-SLC1A2 (1:1000, Abcam, ab41621), anti-FH (1:1000, ProteinTech, 11375-1-AP), anti-PSMB7 (1:1000, Cell Signaling Technology, 13207), anti-UBE4B (1:1000, Abcam, ab126759), anit-MARCH9 (1:1000, Invitrogen, PA5-103817), anti-Skp1 (1:1000, Cell Signaling Technology, 2156), and anti-UBA52 (1:1000, Abcam, ab109227). After two washes for 5 min with 1X wash buffer A, PLA probe solution was applied for 1 h at 37 °C. Tissues were washed and incubated with ligation buffer for 30 min at 37 °C followed by washing and adding the amplification solution for 100 min at 37 °C. Slides were then washed 10 min twice with 1X washing buffer B at RT before mounting using Duolink *In Situ* Mounting Medium with DAPI.

### Confocal image analysis

Brain sections were imaged with UIS2XLine Plan Apochromat (UPLAPO) 63×oil objective (NA 1.5, Olympus) and a 3.2× MagChanger of an Olympus SpinSR-10 Yokogawa spinning disk confocal through an ORCA Fusion sCMOS camera (Hamamatsu). To build the z-stack, 17 stacks/0.37–0.41-μm optimal thickness was captured. Each brain section was randomly imaged in 3 different regions of interest [61, 68]. Laser power was set using secondary antibody–only controls. All images were analyzed using ImageJ. Each image was converted into 8-bit greyscale for each channel. The channel containing the PLA signal was thresholded until the PLA puncta were reliably isolated from the background forming a binary image. Overlapping PLA puncta were segmented using a watershed function and over/undersized puncta were excluded by defining the expected size range of PLA puncta. the summary result of PLA puncta/field was reported [30, 73].

### Western blot and dot blot analysis

For Western blot analysis, an equal proportion of PBS- or SRK-soluble fractions from human brain homogenate was loaded on precast NuPAGE 4 to 12% Bis-Tris gels (Invitrogen) for SDS-PAGE analysis. Gels were subsequently transferred onto nitrocellulose membranes and blocked for 1 h at RT using Odyssey Blocking Buffer (LI-COR). For dot blot, an equal amount of IP-T18 samples was dotted on the membrane and let dry for 1 h at RT followed by blocking. Membranes were then probed overnight at 4 °C using the corresponding primary antibodies followed by IRdye secondaries (LI-COR) at 1:10,000 for 1 h at RT. Images were acquired by LI-COR Odyssey imager. Densitometric analysis was performed using ImageJ software (NIH). The primary antibodies used were the following: mouse anti-Tau 13 (1:20000, BioLegend; MMS-520R), rabbit anti-T18 (1:2000), rabbit anti-FABP5 (1:2000, ProteinTech, 12348-1-1AP), rabbit anti-SLC1A2 (1:1000, Abcam, ab41621), rabbit anti-FH (1:1000, ProteinTech, 11375-1-AP), rabbit anti-PSMB7 (1:1000, Cell Signaling Technology, 13207), rabbit anti-UBE4B (1:1000, Abcam, ab126759), rabbit anit-MARCH9 (1:1000, Invitrogen, PA5-103817), rabbit anti-Skp1 (1:1000, Cell Signaling Technology, 2156), rabbit anti-UBA52 (1:1000, Abcam, ab109227), mouse anti-MARS (1:2000, Abcam, ab50793), rabbit anti-MAGT1 (1:1000, ProteinTech, 17430-1-AP), and rabbit anti-GAPDH antibody (1:1000, Abcam; ab9485).

### Mass spectrometry

**Regents and chemicals** – all reagents were ACS grade or higher. All solvents used, including water, were LC/MS grade. Ammonium bicarbonate (ABC), 2,2,2-trifluoroethanol (TFE), and acetic acid were purchased from Sigma-Aldrich. Iodoacetamide (IDA), dithiothreitol (DTT), acetonitrile (ACN), formic acid, and methanol were purchased from Thermo Scientific. Urea ultra was from MP Biomedicals. Sequencing-grade modified trypsin (Promega) was used.

### Trypsin digestion

The trypsin digestion was performed as previously described [82, 95]. 100 µL of 50 mM ammonium bicarbonate was added to each sample. The beads were suspended with gentle vortexing for 1h. The proteins on the beads were reduced with 10 mM DTT for 30 min, then alkylated with 20 mM IDA for 1h in dark. An aliquot of 4 µg of sequencing-grade trypsin was added to each sample before a 4 h incubation at 37°C with gentle shaking; the supernatant was then collected. Another 4 µg of trypsin was then added to the beads, and the sample was incubated at 37°C overnight with gentle shaking; the supernatant was then collected. After trypsin digestion, the beads were washed twice with 50 µL of 50% ACN, and the supernatants were collected. All of the supernatants were combined and dried with a SpeedVac.

### LC-MS/MS analysis

A nanoflow ultra-high performance liquid chromatography (UHPLC) instrument (Easy nLC, Thermo Fisher Scientific) was coupled on-line to a Q Exactive mass spectrometer (Thermo Fisher Scientific) with a nanoelectrospray ion source (Thermo Fisher Scientific). Peptides were loaded onto a C18-reversed phase column (25 cm long, 75 μm inner diameter) and separated with a linear gradient of 5–35% buffer B (100% acetonitrile in 0.1% formic acid) at a flow rate of 300 nL/min over 120 min. Each sample was analyzed by LC-MS/MS twice. MS data were acquired using a data-dependent Top15 method, dynamically choosing the most abundant precursor ions from the survey scan (350–1600 m/z) using HCD fragmentation. Survey scans were acquired at a resolution of 70,000 at m/z 400. Unassigned precursor ion charge states as well as singly charged species were excluded from fragmentation. The isolation window was set to 3 Da and fragmented with normalized collision energies of 28. The maximum ion injection times for the survey scan and the MS/MS scans were 20 ms and 120 ms respectively, and the ion target values were set to 3E6 and 1e5, respectively. Selected sequenced ions were dynamically excluded for 30 seconds. Data was acquired using Xcalibur software.

### Data processing and Bioinformatic Analysis

Raw MS data were analyzed using MaxQuant software version 1.5.2.8 using the Andromeda search engine [17, 18]. The initial maximum allowed mass deviation was set to 10 ppm for monoisotopic precursor ions and 20 ppm for MS/MS peaks. Enzyme specificity was set to trypsin, defined as C-terminal to arginine and lysine excluding proline, and a maximum of two missed cleavages was allowed. Carbamidomethylcysteine was set as a fixed modification, methionine oxidation, ubiquitination of lysine, acetylation of lysine, phosphorylation of serine, threonine, and tyrosine as variable modifications. The spectra were searched with the Andromeda search engine against the SWISSPROT human sequence database (containing 42,130 human protein entries), combined with 248 common contaminants and concatenated with the reversed versions of all sequences. Protein identification required at least one unique or razor peptide per protein group. Quantification in MaxQuant was performed using the built-in XIC-based label-free quantification (LFQ) algorithm [17]. The required false positive rate for identification was set to 1% at both the peptide and protein level, and the minimum required peptide length was set to 8 amino acids.

For statistical analysis of MaxQuant output we used the Perseus platform (Version 1.6.15.0) [85]. Contaminants, reverse identification, and proteins only identified by site were excluded from further data analysis. The LFQ values were log_2_-transformed. After removing the proteins with less than three valid LFQ values in total, the remaining missing LFQ values were imputed from a normal distribution (width, 0.3; down shift, 1.8).

### Network analysis and visualization

The significant proteins were imported to Cytoscape (Version 3.10.1) [45] for network analysis and annotation enrichment analysis. The visualization and functional enrichment analysis of the protein- protein interaction network of soluble tau was conducted using Cytoscape StringApp [22].

Gene Ontology term enrichment analyses were performed using EnrichmentMap tool [72]. Enrichment and network analyses were applied from Gene Ontology Biological Process (GOBP), Kyoto Encyclopedia of Genes and Genomes (KEGG), and Reactome. Benjamini-Hochberg false discovery rate below 5% was used as the cutoff for significant terms.

### Parallel reaction monitoring-mass spectrometry (PRM) Analysis of ubiquitination of tau

For PRM analyses, the peptides were analyzed with Easy nLC1000 UHPLC-Q Exactive Orbitrap LC-MS system (Thermo Scientific, San Jose, CA). The resolution of full scan was 70,000 (@m/z 200), the target automatic gain control value was set to 3 × 10^6^, and maximum fill time was 200 milliseconds for full scan; and 17,500 (@m/z 200), a target automatic gain control value of 2 × 10^5^, and maximum fill times of 100 milliseconds for MS2 scan. PRM targeted ubiquitinated tau peptides. The assessment of the detection of peptides was performed after acquisition using Skyline version 3.6 [81]. For each peptide evaluated, the signals of the 3-5 most intense fragment ions were extracted from each corresponding MS/MS spectrum library of these ubiquitinated tau peptides. The MS/MS spectra with five fragment ions detected were submitted to spectral matching. The comparison of the relative intensities of these fragments with those defined in the reference composite MS/MS spectrum was performed on the basis of the dot product (dotp) value.

### Statistical analysis

Statistical analyses were performed using Prism 10.0 (GraphPad Software) through unpaired two-tailed Student’s *t* test or one-way analysis of variance (ANOVA) according to group number. Results are considered statistically significant at *p* < 0.05.

## DATA AVAILABILITY

All data are contained within this manuscript and supporting information. The datasets used during the current study are available from the corresponding authors upon request. Schematic diagrams in Fig. 1A, 1B, 6A, 7A, 8, Sup. Fig. 1B, 1J, and 5A were generated with BioRender.

## AUTHOR CONTRIBUTIONS

N.P., Y.Z., and R.K. conceptualized the study; N.P., Y.Z., and R.K. developed the methodological approaches; R.K. and Y.Z. provided the lab resources; N.P., N.B., N.S., E.M., M.S., A.B., J.L., and Y.Z. conducted the experiments; N.P., N.B., and Y.Z. analyzed the data; N.P., NB, M.M., Y.Z., and R.K. wrote the original draft; All authors contributed to reviewing and editing the manuscript, and approved its final version.

## ACKNOWLEDGEMENTS

The authors acknowledge the technical and scientific assistance of Dr. Yu-Hsiu Wang and the Biochemistry and Molecular Biology confocal microscopy core at the University of Texas Medical Branch. We thank the members of Dr. Kayed’s lab for their help and support.

## FIGURE LEGENDS

**Supplementary Figure 1, related to Figure 1.**
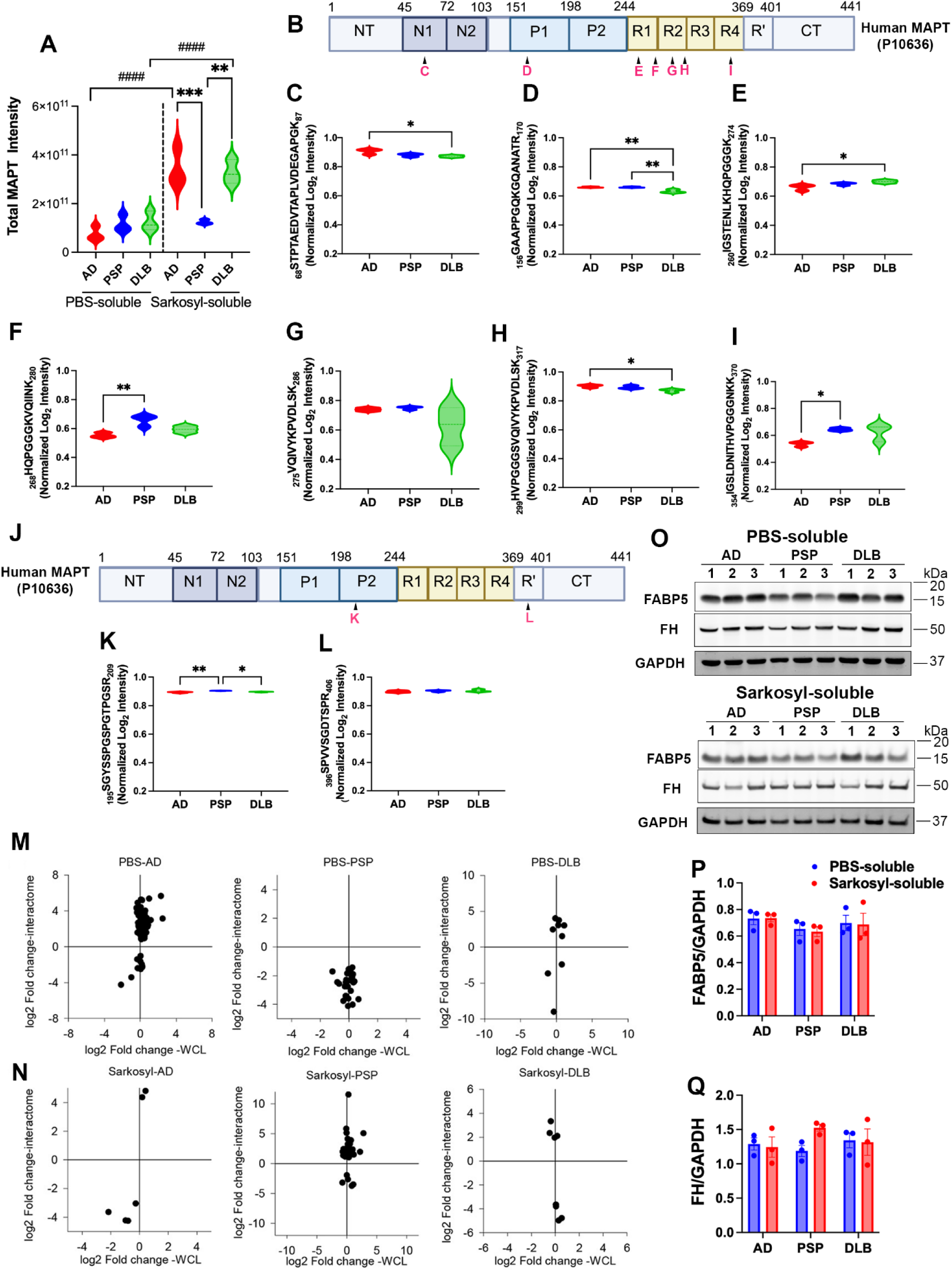
Comparison of misfolded tau interactomes with brain proteome and validation of selected proteins. **(A)** Total MAPT signal intensity across PBS-soluble and sarkosyl-soluble fractions in AD (red), PSP (blue), and DLB (green) cases (n= 3) was detected by label-free LC-MS/MS. **(B)** Domain map of the full-length human MAPT with magenta letters marking the positions of representative tryptic peptides plotted for misfolded tau from PBS soluble fraction (C-I) **(C–I)** Normalized Log2 LFQ intensities for representative tau peptides from PBS-soluble fractions across AD, PSP, and DLB. (**J**) MAPT domain map with peptide positions for misfolded tau from sarkosyl-soluble fraction (**K** and **L**). (**K–L**) Normalized Log2 LFQ intensities for representative tau peptides from sarkosyl-soluble fractions across AD, PSP, and DLB. **(M)** Scatter plots comparing Log2 fold change (FC) in the misfolded tau interactome (y-axis) versus Log2 FC in total PBS soluble lysate (x-axis) for PBS-soluble fractions from AD, PSP, and DLB brains. Each point is a protein significantly detected in T18 immunoprecipitations; grey crosshairs mark zero on both axes. **(N)** Scatter plots comparing Log2 FC in the misfolded tau interactome (y-axis) versus Log2 FC in total sarkosyl-soluble lysate (x-axis) for sarkosyl-soluble fractions from AD, PSP, and DLB brains. Each point is a protein significantly detected in T18 immunoprecipitations; grey crosshairs mark zero on both axes. **(O-Q)** Representative immunoblots and quantification of FABP5 and FH normalized to GAPDH in total PBS-soluble (top) and sarkosyl-soluble (bottom) from AD, PSP, and DLB cases (n=3). Blue dots, PBS- soluble; red dots, sarkosyl-soluble. Quantification of band intensity was normalized to GAPDH. The same immunoblots probed with loading controls shown in Fig. 6. and **Sup. Fig. 3**. Statistical analyses were calculated by Two-way ANOVA with Tukey test. Asterisks indicate pairwise differences among diseases within a fraction; hash marks indicate PBS- vs sarkosyl-soluble differences within the same disease (* p<0.05, ** p<0.01, *** p<0.001, **** p<0.0001; # p<0.05, ## p<0.01, ### p<0.001, #### p<0.0001).

**Supplementary Figure 2, Related to Figure 2.**
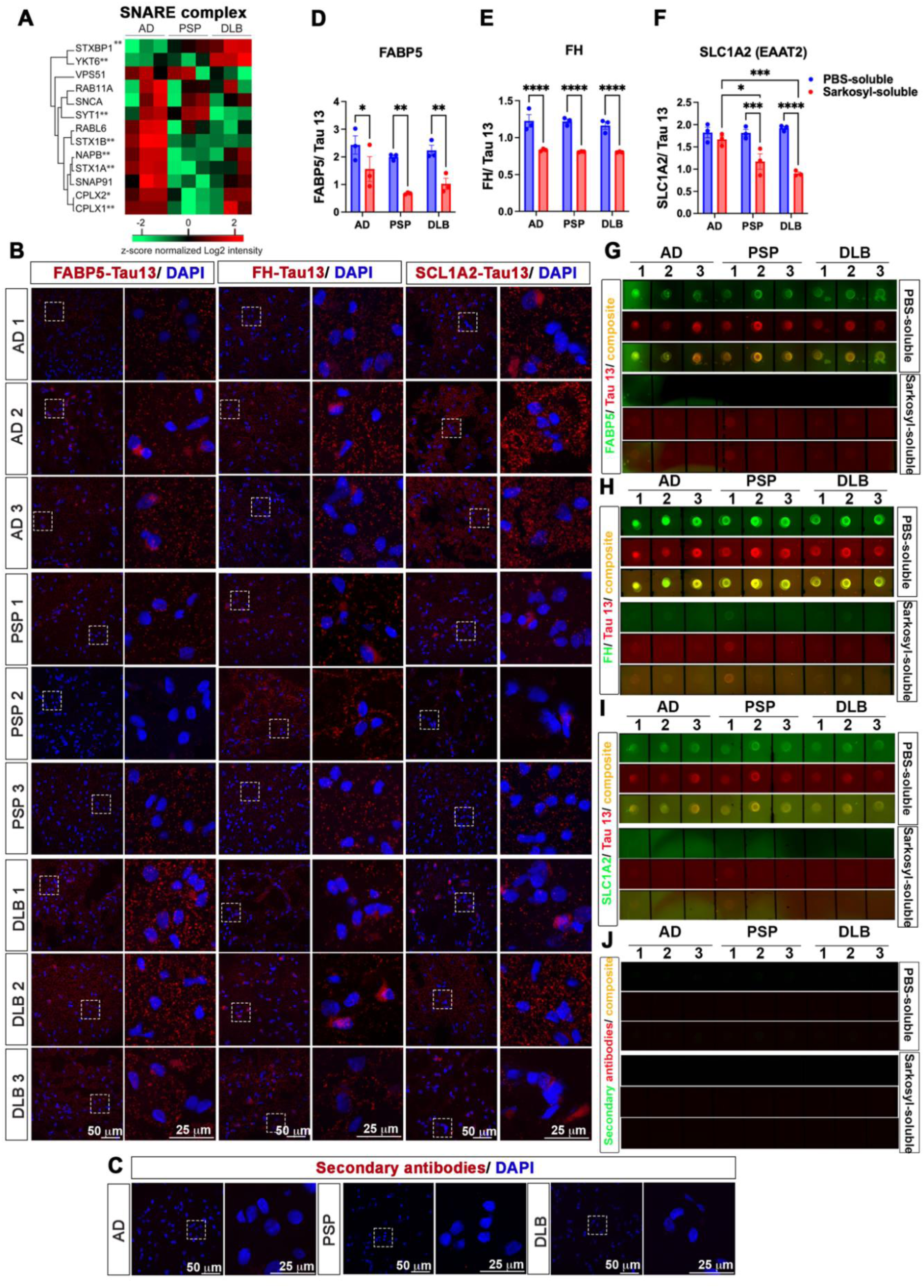
**(A)** SNARE complex interacts with disease-relevant misfolded tau. Heatmap of proteins involved in SNARE complex. **, Class A interactor (q<0.01); *, Class B interactor (q<0.05). Interactome analysis identified 13 SNARE complex proteins, with 9 showing disease-specific interaction patterns. STXBP1 and YKT6 were enriched with DLB derived tau, while SYT1 was depleted from DLB derived tau. Several SNARE proteins, including STX1A, STX1B, and NAPB, significantly interacted with AD tau, but no significant SNARE enrichment was observed with PSP derived tau. CPLX1 and CPLX2 were significantly depleted from PSP derived tau compared to AD- or DLB derived tau. These results highlight disease-specific tau aggregate interactions with SNARE components involved in synaptic vesicle function. **(B)** PLA images showing Tau13 (total tau) in proximity with FABP5, FH, and SLC1A2 in postmortem cortex from AD (AD1–3), PSP (PSP1–3), and DLB (DLB1–3). Boxed regions are magnified at right. Scale bars: 50 μm (left), 25 μm (right). **(C)** Control immunostaining for secondary antibodies only shows no background signal across disease groups (AD, PSP, DLB). (**D–J**) Dual-color dot blot assays of IP materials with T18 antibody (misfolded tau) from PBS- and sarkosyl-soluble fractions probed with Tau13 and antibodies against (**D, G**) FABP5, (**E, H**) FH, and (**F, I**) SLC1A2 in AD, PSP, and DLB samples. Blue dots, PBS-soluble; red dots, sarkosyl-soluble. Data presented as mean ± SEM. *p < 0.05, **p < 0.01, ****p < 0.0001 by one-way ANOVA with Tukey test. **(J)** Dot blot negative control (secondary antibodies only) for PBS- and sarkosyl-soluble IP materials with T18 antibody from AD, PSP, and DLB showing no detectable signal.

**Supplementary Figure 3, related to Table 2.**
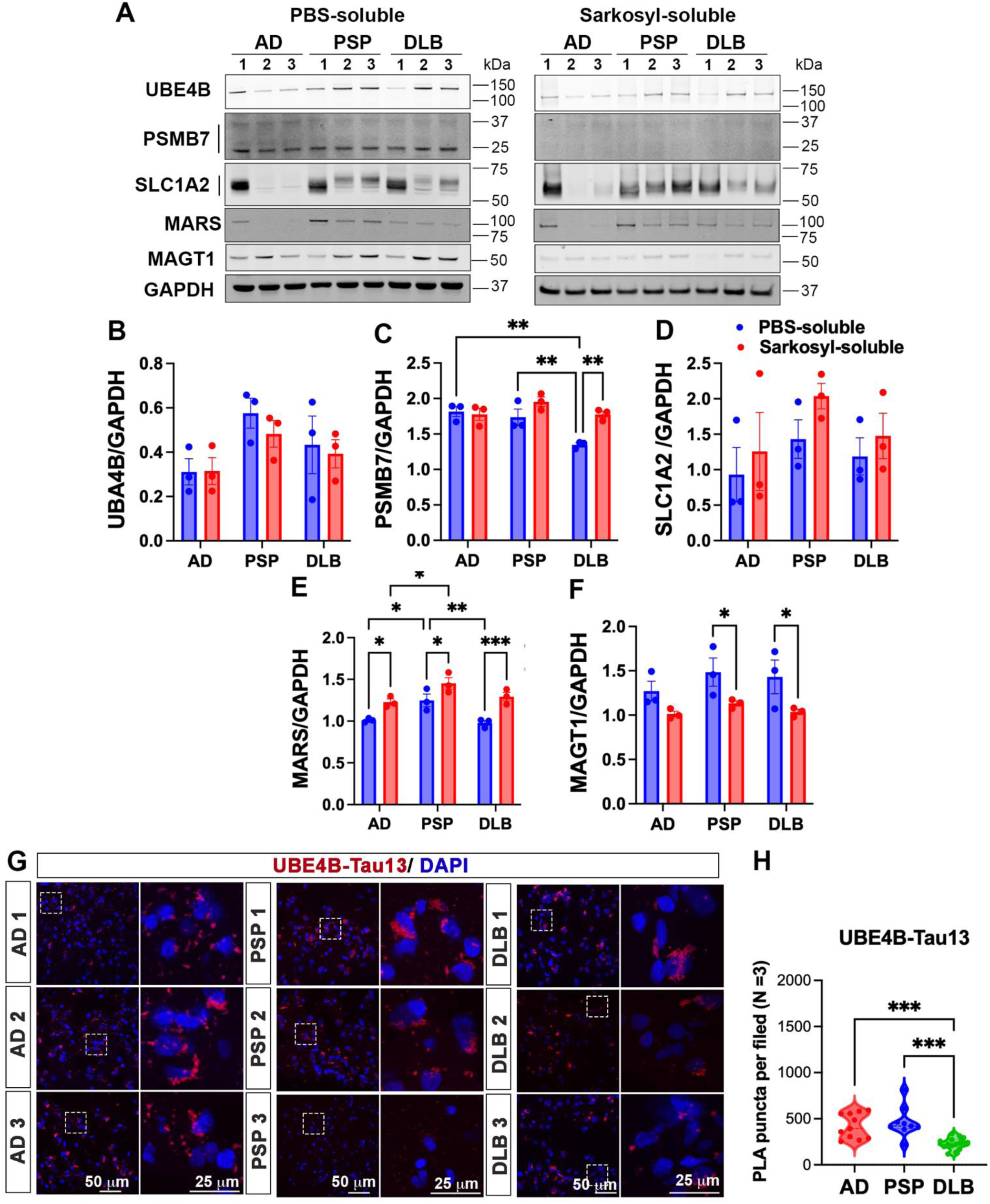
Validation of top-ranked misfolded-tau interactors from PBS- and Sarkosyl-soluble brain fractions. **(A-F)** Representative immunoblots (**A**) and densitometric quantification of immunoblots, normalized to GAPDH of (**B**) UBE4B, (**C**) PSMB7, (**D**) SLC1A2, (**E**) MARS and (**F**) MAGT1 in PBS-soluble and Sarkosyl-soluble fractions from AD, PSP, and DLB (n = 3 per group). Quantification of band intensity was normalized to GAPDH. The same immunoblots probed with loading controls shown in Fig. 6. and **Sup. Fig. 1.** Data presented as mean ± SEM. *p < 0.05, **p < 0.01, ***p < 0.001 by one-way ANOVA with Tukey test. (**G**) PLA images showing colocalization of Tau13 with UBE4B across AD, PSP, and DLB brain samples. Scale bars: 50 μm (left panels), 25 μm (zoom, right panels). (**H**) Quantification of PLA puncta per field for UBE4B–Tau13 colocalization across disease groups (n = 3). Data presented as mean ± SEM. ***p < 0.001 by one-way ANOVA with Tukey test.

**Supplementary Figure 4, related to Figure 6.**
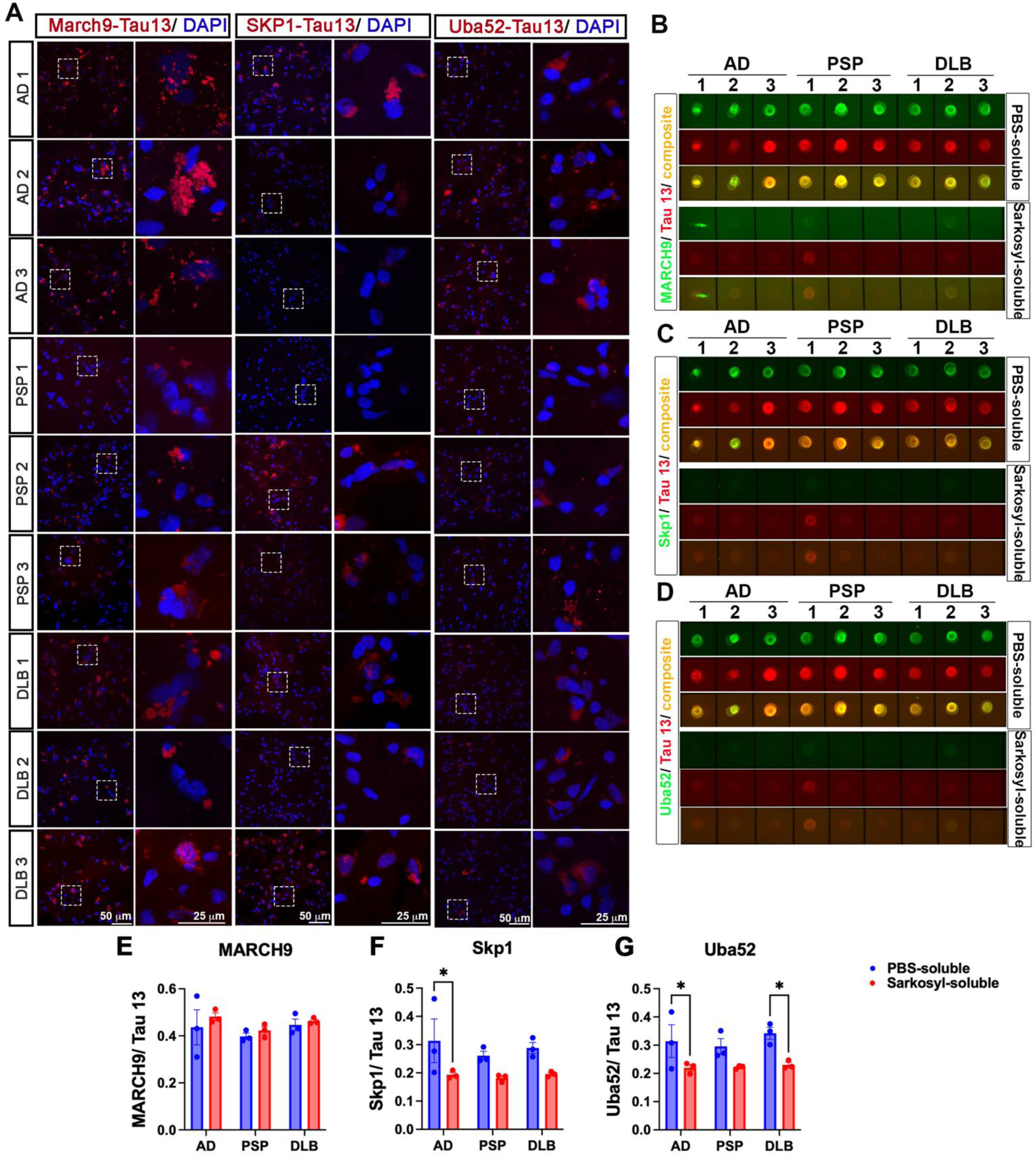
(**A**) Representative PLA images showing colocalization of Tau13 with ubiquitin ligase MARCH9, ubiquitin ligase complex Skp1, and ubiquitin-dependent protein degradation UBA52 in postmortem brain tissue from AD (AD1–3), PSP (PSP1–3), and DLB (DLB1–3) cases Boxed regions are magnified at right. Scale bars: 50 μm (left panels), 25 μm (zoom, right panels). (**B-D**) Dot blot analysis of tau immunoprecipitated from PBS- and sarkosyl-soluble brain fractions using Tau13 in combination with antibodies against (**B**) MARCH9, (**C**) Skp1, or (**D**) UBA52. Tau was immunoprecipitated using the T18 antibody from AD, PSP, and DLB PBS-and sarkosyl-soluble fractions (n = 3 per disease group). (**E–G**) Quantification of dot-blot signals expressed as protein interactor/Tau13 intensity ratios for (**E**) MARCH9, (**F**) SKP1, and (**G**) UBA52 across AD, PSP, and DLB in PBS (blue) and Sarkosyl (red) fractions. Data are mean ± SEM (n = 3 per disease); statistics by two-way ANOVA with Tukey’s post hoc test (*p < 0.05, **p < 0.01).

**Supplemental Figure 5, related to Figures 6 and 7.**
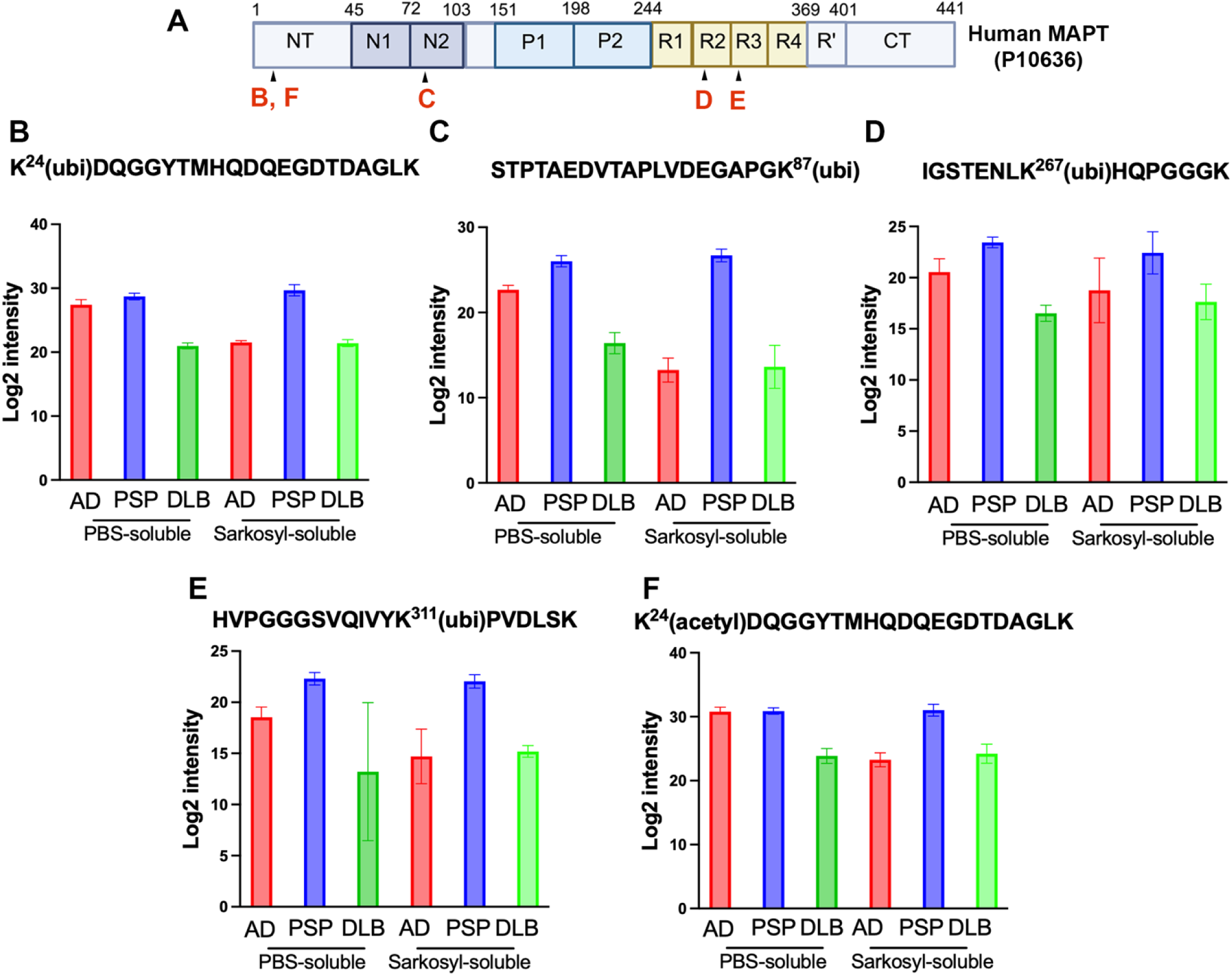
PRM–MS validation of tau ubiquitination and acetylation in the misfolded-tau interactome. **(A)** Domain map of human MAPT (UniProt P10636) with red letters mark the locations of the quantified peptides. (**B–E**) Targeted PRM–MS quantification of four ubiquitinated tau peptides in T18 immunoprecipitates from PBS-soluble and Sarkosyl-soluble fractions of AD, PSP, and DLB brains (n = 3 per disease). Sequences shown on each panel with modified lysines marked “[ubi]”. Bars are log₂ PRM intensities (mean ± SEM). (**E**) Targeted PRM–MS quantification of an acetylated tau peptide (K^24^[ac]DQGGYTMHQDQEGDTDAGLK) across the same samples and fractions. Bars are log₂ PRM intensities (mean ± SEM).

